# Global reinforcement of DNA methylation through enhancement of RNA-directed DNA methylation ensures sexual reproduction in rice

**DOI:** 10.1101/2020.07.02.185371

**Authors:** Lili Wang, Longjun Zeng, Kezhi Zheng, Tianxin Zhu, Yumeng Yin, Dachao Xu, Huadong Zhan, Yufeng Wu, Dong-Lei Yang

**Affiliations:** State Key Laboratory of Crop Genetics and Germplasm Enhancement, Nanjing Agricultural University, 210095, Nanjing, China

## Abstract

DNA methylation is an important epigenetic mark that regulates the expression of genes and transposons. RNA-directed DNA methylation (RdDM) is the main molecular pathway responsible for *de novo* DNA methylation in plants. In Arabidopsis, however, mutations in RdDM genes cause no visible developmental defects, which raising the question of the biological significance of RdDM in plant development. Here, we isolated and cloned *Five Elements Mountain 1* (*FEM1*), which encodes an RNA-dependent RNA polymerase. Mutation in *FEM1* substantially decreased genome-wide CHH methylation levels and abolished the accumulation of 24-nt small interfering RNAs. Moreover, male and female reproductive development was disturbed, which led to the sterility of *fem1* mutants. In wild-type (WT) plants but not in *fem1* mutants, genome-wide CHH DNA methylation levels were greater in panicles, stamens, and pistils than in seedlings. The global increase of methylation in reproductive organs of the WT was attributed to enhancement of RdDM activity including FEM1 activity. More than half of all encoding genes in the rice genome overlapped with hypermethylated regions in the sexual organs of the WT, and many of them appear to be directly regulated by an increase in DNA methylation.

Our results demonstrate that a global increase of DNA methylation through enhancement of RdDM activity in reproductive organs ensures sexual reproduction of rice.

## INTRODUCTION

DNA methylation is a stable epigenetic mark that represses transposon expression and regulates gene expression in eukaryotes. In animals, DNA methylation undergoes reprogramming during development, and an error in DNA methylation or DNA demethylation can cause severe developmental defects (Smith and Meissner, 2013; Greenberg and Bourc’His, 2019). Although DNA methylation in mammals mainly involves CG sequence contexts, DNA methylation in plants involves three sequence contexts: CG, CHG, and CHH (H = A, C, or T) (Law and Jacobsen, 2010; Du et al., 2015; Zhang et al., 2018). The orthologue of DNA methyltransferase 1 (DNMT1), METHYLTRANSFERASE 1 (MET1), works together with three homologous proteins, VARIANT IN METHYLATION 1 (VIM1), VIM2, and VIM3, to maintain CG methylation during DNA replication (Kankel et al., 2003; Saze et al., 2003; Woo et al., 2007; Stroud et al., 2013). CHROMOMETHYLASE 3 (CMT3) is responsible for the maintenance of CHG methylation by recognizing the dimethylated state of histone H3 lysine 9 (H3K9me2), which is established by KRYPTONITE/SUPPRESSOR OF VARIEGATION 3-9 HOMOLOGUE PROTEIN 4 (KYP/SUVH4), SUVH5, and SUVH6 (Bartee et al., 2001; Lindroth et al., 2001; Du et al., 2012). Maintaining CHH methylation during DNA replication is not possible because proteins reading asymmetric sequences are lacking. As a consequence, the maintenance of CHH methylation depends on *de novo* methylation.

There are two molecular pathways for *de novo* methylation of CHH, CG, and CHG in plants. In one pathway, a chromatin-remodeling protein, DECREASE IN DNA METHYLATION 1 (DDM1), releases Histone 1 (H1) to allow CHROMOMETHYLASE 2 (CMT2) to catalyze DNA methylation on long transposons (Zemach et al., 2013; Stroud et al., 2014). Another pathway is called RNA-directed DNA methylation (RdDM), which involves the biosynthesis of small non-coding RNAs and long non-coding RNAs (Zhang and Zhu, 2011; Matzke and Mosher, 2014; Matzke et al., 2015). Facilitated by four homologous chromatin-remodeling proteins (CLSY1-CLSY4) and a transcription factor-like protein DNA-BINDING TRANSCRIPTION FACTOR 1/SAWADEE HOMEODOMAIN HOMOLOG 1 (DTF1/SHH1) (Smith et al., 2007; Law et al., 2013; Zhang et al., 2013; Yang et al., 2018; Zhou et al., 2018), RNA polymerase IV (Pol IV) and RNA-dependent RNA polymerase 2 (RDR2) bind the targets and work together to synthesize Pol IV-dependent small RNAs (P4-RNAs) consisting of 25 to 50 nt (Blevins et al., 2015; Zhai et al., 2015; Yang et al., 2016; Ye et al., 2016). P4-RNAs are cleaved by DICER-LIKE 3 (DCL3) to produce 24-nt siRNAs (Xie et al., 2004; Henderson et al., 2006), which are loaded into ARGONAUTE 4 (AGO4) and ARGONAUTE 6 (AGO6) (Mi et al., 2008; Duan et al., 2015). The long non-coding RNAs transcribed by polymerase V (Pol V) then pair with siRNAs in AGO4 (Wierzbicki et al., 2009). During this process, AGO4 recruits DOMAINS REARRANGED METHYLTRANSFERASE 2 (DRM2) to the DNA targets and catalyzes DNA methylation (Zilberman et al., 2003; Zhong et al., 2014).

The mechanism of RdDM is well-understood in Arabidopsis, but because most Arabidopsis RdDM mutants lack visible developmental defects (Matzke and Mosher, 2014), and because only small numbers of transposons are released in Arabidopsis RdDM mutants (Zemach et al., 2013), the biological role of RdDM is unclear. Rice might be a better model than Arabidopsis for studying the biological role of RdDM in plants because rice has a much larger genome (466 vs. 125 Mb) and a much higher percentage of transposable elements (TEs, 39.5 vs. 18.5%) (Huang et al., 2012; Ausin et al., 2016); the size of the genome and the percentage of transposable elements in rice are more representative of plants in general than is the case for Arabidopsis. There is also evidence that *de novo* methylation affects development in rice. *OsDCL3a* knock-down rice plants, for example, exhibit dwarfism and a larger flag leaf angle (Wei et al., 2014). Mutation in *OsNRPD1* causes increased tillering via its regulation of *OsMIR156* and *DWARF 14* (*D14*) (Xu et al., 2020). *OsDRM2* mutants caused pleiotropic developmental defects at vegetative and reproductive stages (Moritoh et al., 2012; Tan et al., 2016). In tomato, which has a 900 Mb genome (Tomato Genome Consortium, 2012), mutations in *SlNRPD1* or *SlNRPE1* lead to abnormal leaves and flowers, and small fruits (Gouil and Baulcombe, 2016). Mutation in *MOP1* (*MEDIATOR OF PARAMUTATION 1*), an RNA-dependent RNA polymerase gene, caused late flowering and sometimes a short stature and abnormal tassels in maize (Dorweiler et al., 2000). These pleiotropic developmental defects in rice, tomato, and maize mutants suggest that *de novo* methylation might be important in plant species with larger genomes than Arabidopsis.

In the current study of rice, we found that CHH methylation levels were much higher in WT panicles, stamens, and pistils than in WT seedlings. This difference in the methylation levels in reproductive vs. vegetative tissue, however, was absent in the *fem1* mutant, which exhibited reproductive defects. We demonstrate that the increase in DNA methylation in WT panicles, stamens, and pistils depends on *FEM1*, and that the global increase in the DNA methylation in WT sexual organs regulates gene expression in these organs. The results indicate that the upregulated expression of genes that encode RdDM components, including FEM1, is responsible for the increased CHH DNA methylation in WT reproductive organs and is therefore responsible for the proper development of those organs. Finally, the results suggest that the effects of upregulation of RdDM pathway components on reproductive organs should be studied in other plant species.

## Results

### Isolation and map-based cloning of *Five Elements Mountain 1*

We used the CaMV 35S promoter to ectopically express a gibberellin metabolic gene, *OsGA2ox1* (Sakamoto et al., 2004), in the Geng (japonica) rice variety TP309. As expected, most of the transgenic rice plants exhibited gibberellin (GA)-deficient phenotypes including dwarfism, late flowering, and dark green leaves (Supplemental Figures 1A and 1B). A semi-quantitative RT-PCR assay showed that the expression level in the *Os<u>GA</u>2ox1* <u>e</u>ctopic expression line (GAE) was significantly increased (Supplemental Figure 1C). Interestingly, some offspring of the transgenic rice plants restored the developmental defects caused by GA deficiency (Supplemental Figure 1A), and this was true even for plants containing the 35S::*OsGA2ox1* transgene (Supplemental Figure 1B). The expression level was reduced to the WT level in the normal transgenic rice plants (Supplemental Figure 1C), which suggests that gene silencing occurred in those plants. The gene silencing plants were therefore referred as *Os<u>GA</u>2ox1* <u>S</u>ilencing (GAS) plants.

**Figure 1.**
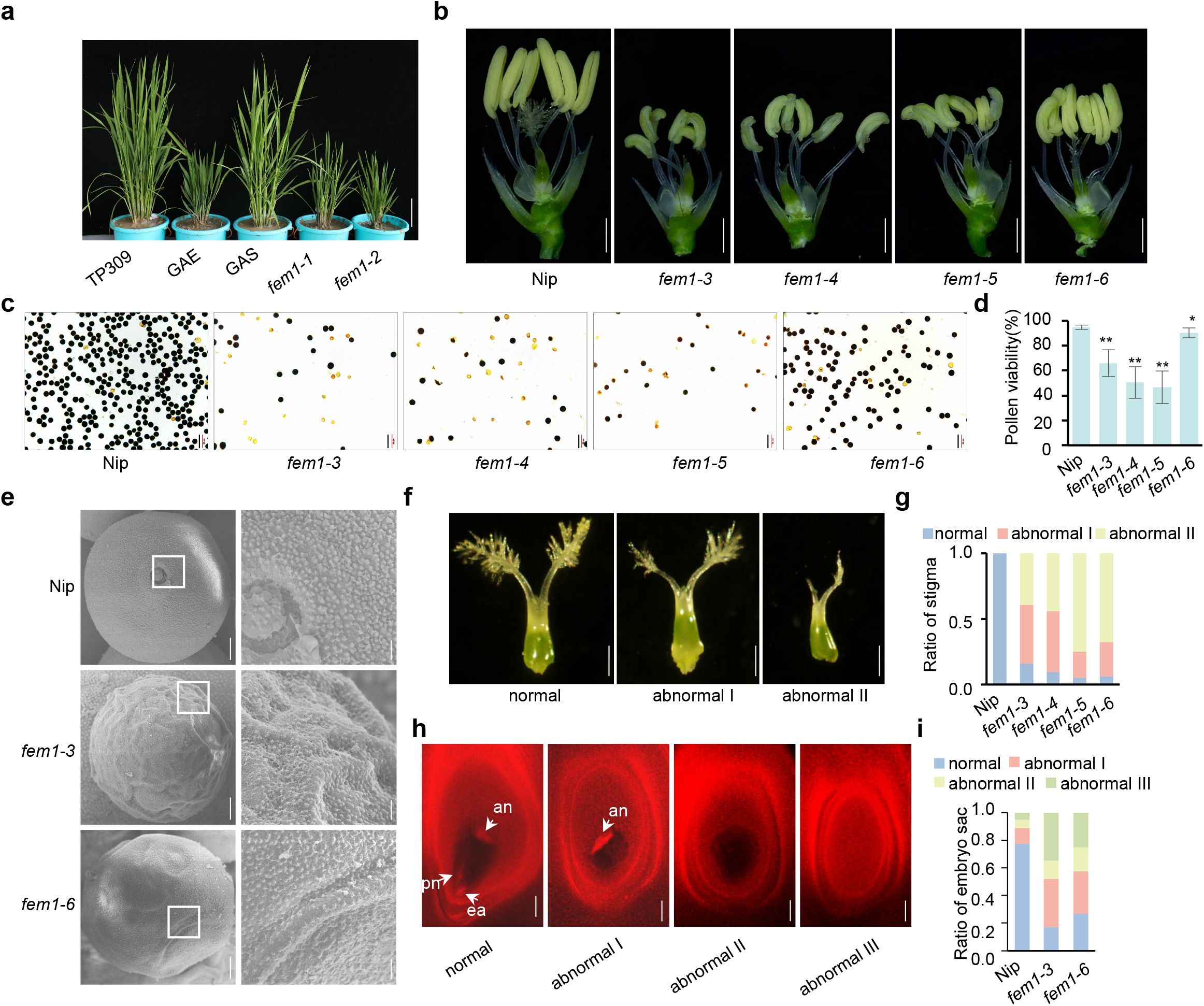
Mutation in *FEM1* causes reproductive defects. (A) Morphology of 3-month-old TP309, GAE, GAS, *fem1-1*, and *fem1-2* plants. Scale bar = 10 cm. (B) Stamen morphology of the WT (Nip) and *fem1* mutants at stage 12. Scale bars = 1 mm. (C) I_2_-KI stained pollen of the WT and *fem1* mutants. Scale bars = 50 μm. (D) Pollen viability as indicated by I_2_-KI staining (**, *p* < 0.01; *, *p* < 0.05, compared to WT by Student’s *t*-test). (E) Scanning electron micrographs of stage 12 pollen grains of the WT and *fem1* mutants. Micrographs on the right are enlargements of those on the left. Scale bars = 5 μm on left, and 1 μm on right. (F) Morphology of mature pistils of normal and abnormal (I and II indicate increasing degrees of abnormality). Scale bars = 0.5 mm. (G) Proportions of stigma phenotypes that are normal and abnormal (I and II) in the indicated genotypes (n = 120). (H) Laser confocal scanning micrographs of normal and abnormal (I, II, and III indicate increasing degrees of abnormality) embryo sacs. an: antipodal cells, pn: polar nucleus, ea: egg apparatus. Scale bars = 100 μm. (I) Proportions of embryo sac phenotypes that are normal and abnormal (I, II, and III) in the indicated genotypes (n = 80).

To elucidate the molecular mechanism of epigenetic silencing of transgenic *OsGA2ox1* in GAS plants, we conducted a genetic screen for genes that suppress gene silencing of 35S::*OsGA2ox1*. More than 23,000 rice plants that germinated from ethyl methyl sulfonate (EMS)-treated GAS seeds were planted and permitted to self-fertilize. The plants with GA-deficient phenotypes like dwarfism and dark green leaves were isolated from the M2 generation. The *OsGA2ox1* expression levels in those plants were then assessed by real-time PCR, and the plants with dramatically increased expression levels (> 20-fold) compared to GAS were further selected and named *five elements mountain* (*fem*) plants (Figure 1A; Supplemental Figures 1D and 1E). We isolated about 200 *fem* mutants, two of which (#45 and #30) were referred to as *fem1-1* and *fem1-2* because subsequent, separate map-based cloning demonstrated that they were allelic to each other (see below).

Because both *fem1-1* and *fem1-2* were totally sterile, we crossed the heterozygous *fem1-1*/*FEM1* (#45) with the Xian (indica) variety TN1 (Taichuang native 1); the dwarf plants from the F2 population were selected and constituted a map-based cloning population. In total, 890 dwarf plants were used to locate the candidate gene in a 103-kb region (Supplemental Figure 1F). Among seven genes in this region, *fem1-1* contained two mutations on LOC_Os04g39160 (both were C to T) that caused two amino acid substitutions near its carboxylic terminal (Supplemental Figure 1F). To confirm the cloning result, we conducted a genetic complementary assay using the native promoter to drive the genomic DNA of LOC_Os04g39160. The transgenic plants restored the GA-deficient phenotypes and reduced *OsGA2ox1* expression (Supplemental Figures 1G to 1I).

Another independent map-based cloning of a different *fem* mutant (#30) also identified one mutation (G to A) in the same gene, a mutation that caused a one-amino acid substitution from glycine to glutamic acid (Supplemental Figure 1F). Moreover, genetic analysis demonstrated that *fem1-1* and *fem1-2* were allelic (Supplemental Figures 1J to 1L). Taken together, these results demonstrated that LOC_Os04g39160 is *FEM1*. A phylogenetic tree showed that FEM1 is homologous with the RDR2 protein in Arabidopsis and with the MOP1 protein in maize (Supplemental Figure 1M).

### Mutation in*FEM1* causes reproductive defects

*fem1-1* and *fem1-2* were probably weak alleles of *FEM1*, and TP309 lacks genomic reference. To overcome these problems, we knocked out *FEM1* using CRISPR/Cas9 technology at two different single-guide RNA (sgRNA) sites of *FEM1* in Nipponbare, which has a high quality reference genome. sgRNA1 targeted the DNA sequence encoding the RNA-dependent RNA polymerase (RdRP) domain, and sgRNA2 targeted the C-terminal (Supplemental Figures 2A and 2B). One-base insertion or deletion near the protospacer-adjacent motif (PAM) in three *fem1* alleles (*fem1-3*, *fem1-4*, and *fem1-5*) generated by sgRNA1 caused a premature stop codon and led to a truncated protein without an intact RdRP domain, indicating that the three *fem1* alleles might be functional null mutants (Supplemental Figure 2B). The *fem1-6* mutant generated by sgRNA2 contained 49 amino acid (aa) mutations with a 12-aa deletion on its C-terminal (Supplemental Figure 2B). Like the *OsDCL3a* and *OsNRPD1* knock-down rice plants (Wei et al., 2014; Xu et al., 2020), *fem1* mutants were dwarfs, and their flag leave angle was much larger than that of WT plants with an empty vector (Supplemental Figure 2C). Interestingly, the anthers of the *fem1* mutants were smaller and paler than those of control plants (Figure 1B). Furthermore, iodine potassium iodide (I_2_-KI) staining showed that the pollen viability was dramatically decreased in *fem1* mutants (Figures 1C and 1D). Electron microscopy showed that the pollen surface was smooth for the WT but wrinkled for the *fem1* mutants (Figure 1E). To better understand the defect in anther development, we analyzed transverse sections of WT and *fem1* anthers. At stage 6, when anther morphogenesis is complete, the WT anthers formed four somatic layers (the epidermis, endothecium, middle layer, and tapetum), and the microsporocytes were located at the center of each anther locule (Supplemental Figure 2D). In *fem1* mutants, however, a portion of the anther lobes was arrested at stage 6 and failed to form the organized sporophytic cell layers, with the exception of the epidermis, indicating that *FEM1* is essential for cell differentiation during early anther development in rice (Supplemental Figure 2D). The anther defects were much weaker in *fem1-6* than in the other three *fem1* mutants (Figures 1B to 1E), indicating a partial loss-of-function in *fem1-6*.

**Figure 2.**
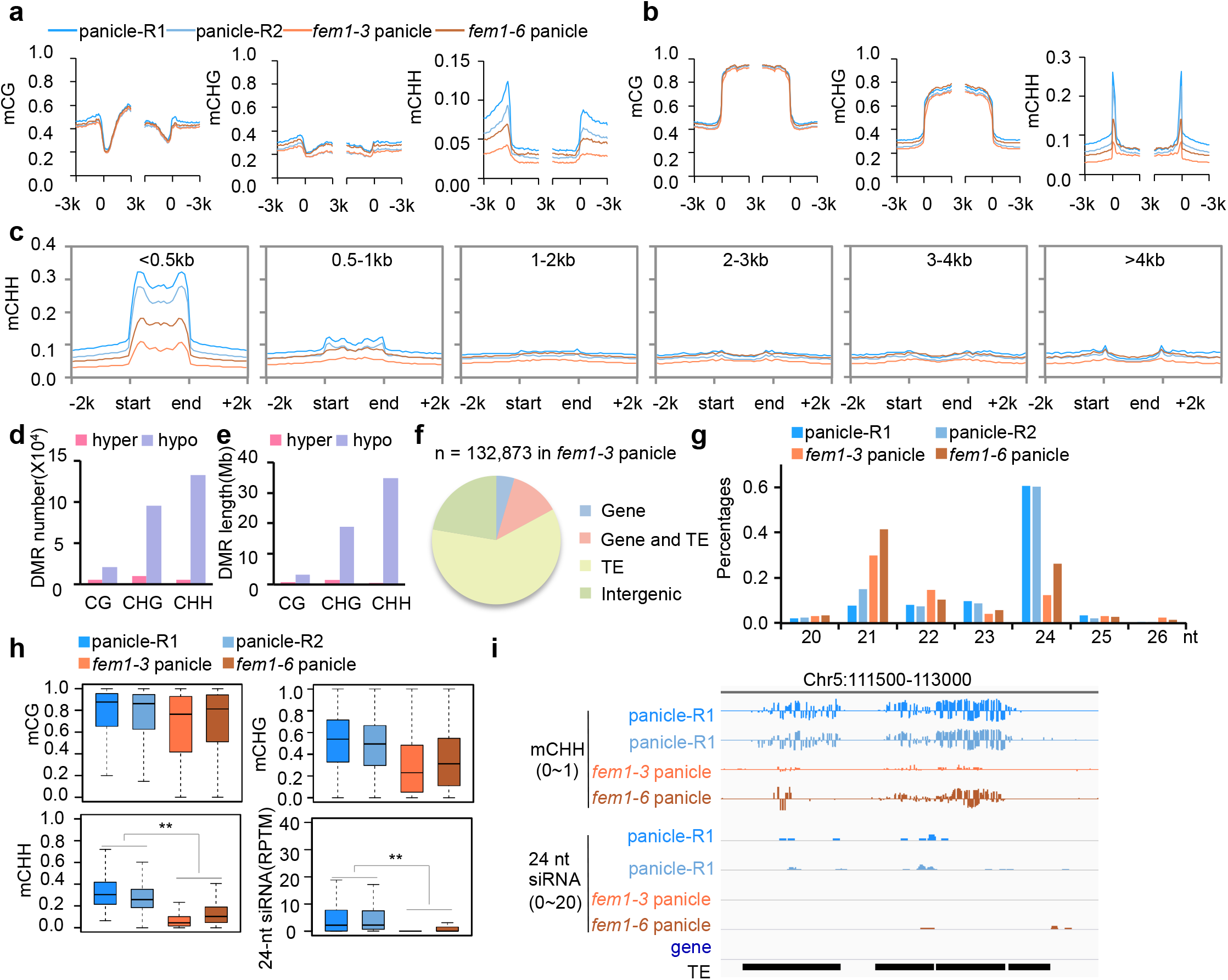
FEM1 regulates CHH methylation and 24-nt siRNA biosynthesis. Normalized CG, CHG, and CHH methylation levels on genes (A) and TEs (B) in the panicles of various genotypes (R1 represent WT-Replicate 1; R2 represent WT-Replicate 2). (C) Normalized CHH methylation levels on TEs of different length in panicles of the indicated genotypes. The number (D) and length (E) of various DMRs in panicles of *fem1-3* relative those in the WT. (F) Genomic distribution of CHH hypo-DMRs in *fem1-3* panicles. (G) Abundance of small RNAs of different length in WT and *fem1* panicles. (H) Box-plots indicating the methylation level of CG, CHG, and CHH, and 24-nt siRNA abundance on CHH hypo-DMR in *fem1-3* panicles (**, *p*<0.01 by Wilcoxon sum test). (I) Integrated Genome Browser view of CHH methylation and 24-nt siRNA abundance in one representative CHH hypo-DMR in panicles of the indicated genotypes.

In addition, stigmas of each gynoecium in *fem1* lacked one or two hairbrushes (Figures 1F and 1G). Abnormal embryo sacs were more frequently found in *fem1* mutants than in the WT (Figures 1H and 1I), indicating that female gametophyte development was disturbed. Thus, mutation in *FEM1* caused both male and female reproductive developmental defects and consequently led to sterility. *fem1-3*, *fem1-4*, and *fem1-5* plants produced no seeds, and *fem1-6* plants produced only a small number of seeds (Supplemental Figures 2E to 2H).

### *FEM1* regulates the methylome and 24-nt siRNAs

To elucidate the role of *FEM1* in the methylome, we conducted whole-genome bisulfite sequencing (WGBS) for the panicles of *fem1-3* and *fem1-6* with the corresponding controls created in the same regeneration process. The high quality of the methylome (Supplemental Data Set 1) showed that mutation in *FEM1* reduces the genome-wide methylation level (GML) of CHH, i.e., the CHH methylation level dropped from 7.0% in WT replicate 1 to 2.9% in *fem1-3*, and from 5.1% in WT replicate 2 to 3.9% in *fem1-6* (Supplemental Data Set 1). For genes, CHH methylation levels were higher for upstream and downstream regions in the WT than for the gene body (Figure 2A). The high CHH methylation levels on the borders of genes were dramatically reduced in *fem1-6* and nearly eliminated in *fem1-3* (Figure 2A), suggesting that CHH methylation upstream and downstream of genes was dependent on *FEM1*. As was the case for genes, CHH methylation but not CG or CHG methylation on TEs was also dependent on *FEM1* (Figure 2B). CHH methylation depends on *FEM1* not only in panicles but also in seedlings (Supplemental Figure 3A and 3B). To determine which kind of TEs were dominant targets of *FEM1*, we divided TEs into six subgroups based on length. On short TEs (< 500 bp), the CHH methylation level was as high as 30% in the WT, but was 10% in the *fem1-3* mutant (Figure 2C). On long TEs (> 500 bp), however, the methylation level in the WT was very low, and the difference between the WT and *fem1* was small (Figure 2C). That *FEM1* mainly regulated DNA methylation on short TEs was also apparent in seedlings (Supplemental Figure 3C).

**Figure 3.**
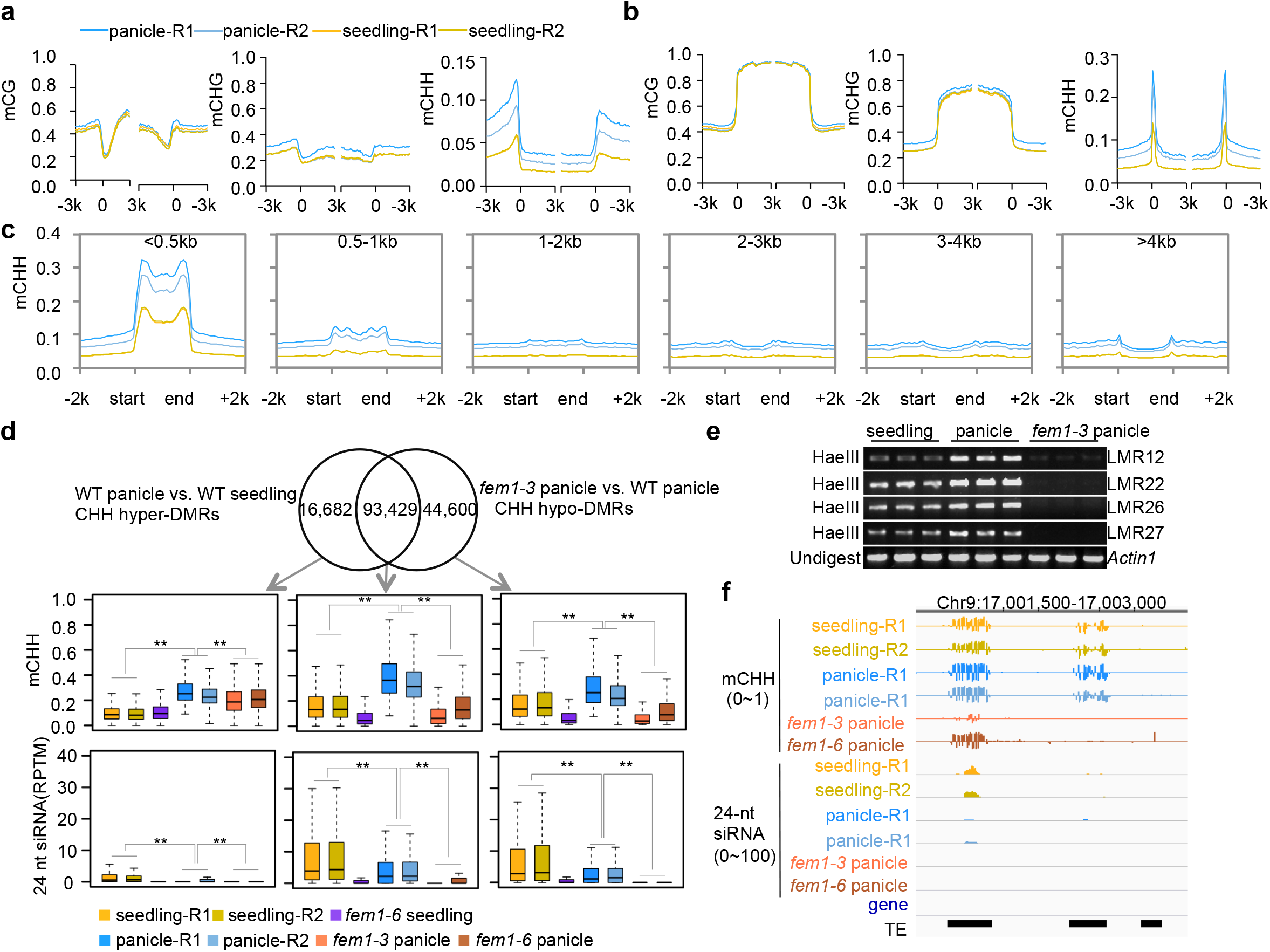
Increase of CHH methylation in panicles depends on *FEM1*. Normalized CG, CHG, and CHH methylation levels on genes (A) and TEs (B) in panicles and seedlings. (C) Normalized CHH methylation levels on TEs of different length in panicles and seedlings. (D) Venn diagram of the overlap between CHH hyper-DMRs of panicles > seedlings and hypo-DMRs of *fem1-3* < WT in panicles. Box-plots indicate the methylation levels of CHH and 24-nt siRNA abundance on common and specific regions in panicles or seedlings of various genotypes (**, *p*<0.01 by Wilcoxon sum test). (E) Chop-PCR assay on several panicle CHH hyper-DMRs. The DNA template was cut by *Hae*III. (F) Integrated Genome Browser view of CHH methylation levels and 24-nt siRNAs abundance in one representative CHH hyper-DMR in panicles and seedlings of the indicated genotypes.

We further identified differentially methylated regions (DMRs) between *fem1* panicles and WT panicles. Many hypo-DMRs but very few hyper-DMRs of CHG and CHH were identified in the *fem1* panicles (Figure 2D; Supplemental Figure 3D; Supplemental Data Set 2), suggesting that *FEM1* positively regulates DNA methylation. Among DMRs in panicles, CHH hypo-DMRs were the longest (Figure 2E; Supplemental Figure 3E). The total length of CHH hypo-DMRs covered 34.8 Mb in *fem1-3* panicles and 23.4 Mb in *fem1-6* panicles (Figure 2E; Supplemental Figure 3E). The majority of CHH hypo-DMRs of *fem1* were located on TEs (Figure 2F; Supplemental Figure 3F). Small RNA sequencing (sRNA-seq) showed that 24-nt siRNAs were greatly reduced in *fem1-6* panicles and nearly eliminated in *fem1-3* panicles (Figure 2G; Supplemental Data Set 1), which was consistent with the conclusion that RDR2 and MOP1 are essential for 24-nt siRNA biosynthesis (Xie et al., 2004; Lu et al., 2006; Nobuta et al., 2008). On the CHH hypo-DMRs of *fem1-3* panicles, 24-nt siRNAs were significantly decreased in *fem1-3* panicles (Figures 2H and 2I), suggesting that the activity of FEM1 is required for both siRNA biosynthesis and DNA methylation.

### Increased DNA methylation in panicles depends on*FEM1*

We also conducted WGBS for *fem1-6* and WT seedlings (Supplemental Data Set 1). As is the case in panicles, the CHH hypo-DMRs are the main type of DMRs in *fem1-6* seedlings (Supplemental Figures 3G and 3H; Supplemental Data Set 2), and the DMRs in seedlings are mainly located on TEs (Supplemental Figure 3I). The number of 24-nt siRNAs in *fem1-6* was greatly decreased globally on the CHH hypo-DMRs in seedlings (Supplemental Figures 3J to 3L), which was consistent with the features in panicles. However, the obvious difference between the number and length of hypo-DMRs in *fem1-6* panicles vs. *fem1-6* seedlings indicated that the DNA methylation level might differ among organs (Supplemental Figures 3D, 3E, 3G and 3H). To understand this difference, we directly compared the GML in the WT panicles and seedlings, and found that CHH methylation levels in panicles were 5.1% and 7.0%, which were substantially higher than the 3.1% in seedlings (Supplemental Data Set 1). On both genes and TEs, the CHH methylation level but not the CG or CHG methylation level was dramatically higher in WT panicles than in WT seedlings (Figures 3A and 3B). The increase in CHH methylation level on transposons mainly occurred on short TEs (Figure 3C). In contrast, the CG and CHG levels in WT panicles and WT seedlings were similar on TEs of various lengths (Supplemental Figures 4A and 4B). CHH hyper-DMRs were the main type of DMRs between panicles and seedlings (Supplemental Figures 4C and 4D, the CHH hyper-DMRs were named “panicle hyper-DMRs”), and these DMRs were also mainly located on TEs (Supplemental Figure 4E). On the panicle hyper-DMRs, the CG methylation level was similar in WT panicles and WT seedlings (Supplemental Figure 4F). The CHH level was substantially higher and the CHG level was significantly higher in WT panicles than in WT seedlings (Supplemental Figure 4F). However, the 24-nt siRNA abundance on the panicle hyper-DMRs was not increased but was significantly lower in WT panicles than in WT seedlings (Supplemental Figure 4F), suggesting that siRNA levels were not responsible for the increase in DNA methylation.

**Figure 4.**
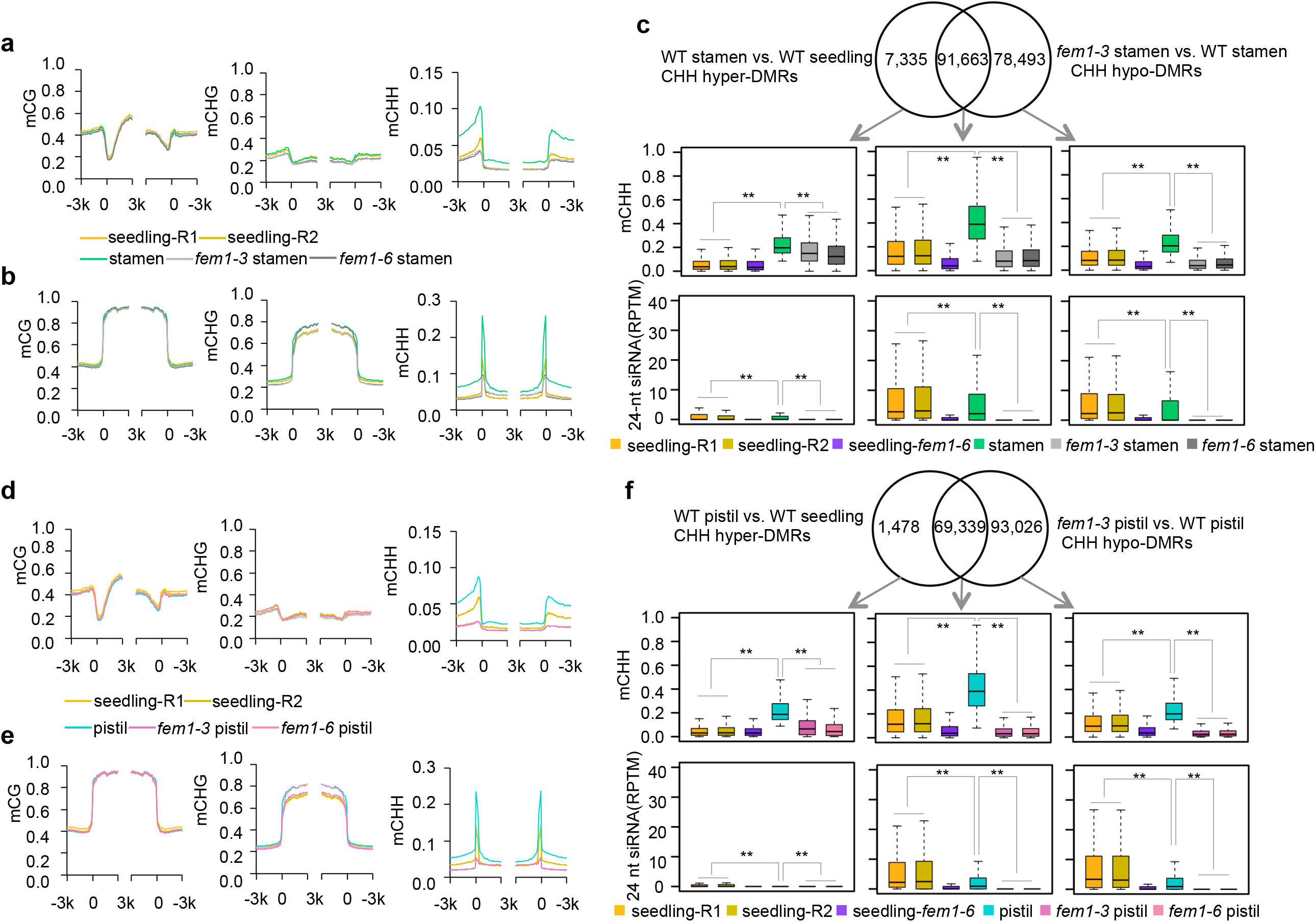
Increase of CHH methylation in stamens and pistils depends on FEM1. Normalized CG, CHG, and CHH methylation levels on genes (A) and TEs (B) in the stamens and seedlings of the indicated genotypes. (C) Venn diagram showing the overlap between stamen CHH hyper-DMRs and *fem1-3* hypo-DMRs in stamens. Box-plots indicate the methylation levels of CHH and 24-nt siRNAs abundance on the overlapped and specific regions in stamens or seedlings of the indicated genotypes (**, *p*<0.01 by Wilcoxon sum test). Comparison of CG, CHG, and CHH methylation levels on genes (D) and TEs (E) in pistils and seedlings of the indicated genotypes. (F) Venn diagram showing the overlap between CHH hyper-DMRs of pistils > seedlings and hypo-DMRs *fem1-3* < WT in pistils. Box-plots indicate the methylation levels of CHH and 24-nt siRNA abundance on the common and specific regions in pistils or seedlings of the indicated genotypes (**, *p*<0.01 by Wilcoxon sum test).

The increased methylation on panicle hyper-DMRs and on the *FEM1*-dependent DMRs in both panicles and seedlings occurred mainly on short TEs and in CHH context, suggesting that the increase methylation in panicles may depend on *FEM1*. Among 108,932 panicle hyper-DMRs, about 85.8% overlapped with *FEM1*-dependent DMRs in the panicle (Figure 3D), which suggested that the majority of panicle hyper-DMRs were *FEM1*-dependent. In the overlapped regions, the high CHH methylation level in WT panicles was reduced in *fem1* panicles (Figure 3D). Even on the specific *FEM1*-dependent DMRs, the CHH methylation level was higher in WT panicles than in WT seedlings, suggesting that the extent of dependence on *FEM1* for increased methylation in panicles was larger. Consistent with that possibility, the CHH methylation level in all *FEM1*-dependent CHH DMRs in panicles was higher in panicles than in seedlings (Supplemental Figure 4G). A Chop-PCR assay and IGV snapshot showed that the increase in CHH methylation level in panicle hyper-DMRs was dependent on *FEM1* (Figures 3E and 3F). However, the increase in CHH methylation was not accompanied by an upregulation of siRNA abundance (Figures 3D and 3F; Supplemental Figures 4F and 4G), which indicated that the change in siRNA abundance did not explain the increase in DNA methylation.

### Increased DNA methylation in stamens and pistils depends on *FEM1*

To determine whether the increased methylation in panicles involves an increase in methylation of the reproductive organs, we performed WGBS of stamens and pistils of the WT, *fem1-3*, and *fem1-6* (Supplemental Data Set 1). On a genome-wide scale, the CHH methylation level was 5.5% in WT stamens and 4.7% in WT pistils, but was only 3.1% in WT seedlings (Supplemental Data Set 1). The CHH methylation level was higher on both genes and TEs in stamens than in seedlings (Figures 4A and 4B). On transposons, CHH methylation level in stamens mainly occurred on short TEs (Supplemental Figure 5A). There were 98,434 CHH hyper-DMRs between stamens and seedlings (Supplemental Data Set 3), which covered 17.8 Mb (Supplemental Figures 5B and 5C), and among the stamen hyper-DMRs, 59.4% were located on TEs (Supplemental Figure 5D). Mutation in *FEM1* produced 162,809 CHH hypo-DMRs in stamens (Supplemental Figure 5E; Supplemental Data Set 3), which covered 45.5 Mb, i.e., about one-tenth of the rice genome (Supplemental Figure 5F), and 58.6% of the CHH DMRs were on TEs (Supplemental Figure 5G). About 93.1% of the stamen hyper-DMRs overlapped with *FEM1*-dependent CHH methylation regions in stamens (Figure 4C). In the overlapped regions, the CHH methylation level was higher in WT stamens than in WT seedlings, and mutation in *FEM1* almost eliminated CHH methylation in both seedlings and stamens (Figure 4C). Like the overlapped DMRs, the CHH levels on the specific *FEM1*-dependent DMRs were higher in WT stamens than in WT seedlings (Figure 4C), suggesting that the increased methylation in stamens was fully *FEM1*-dependent. However, siRNA abundance was not higher in WT stamens than in WT seedlings on those regions (Figure 4C), indicating an inconsistency between siRNA abundance and methylation levels.

**Figure 5.**
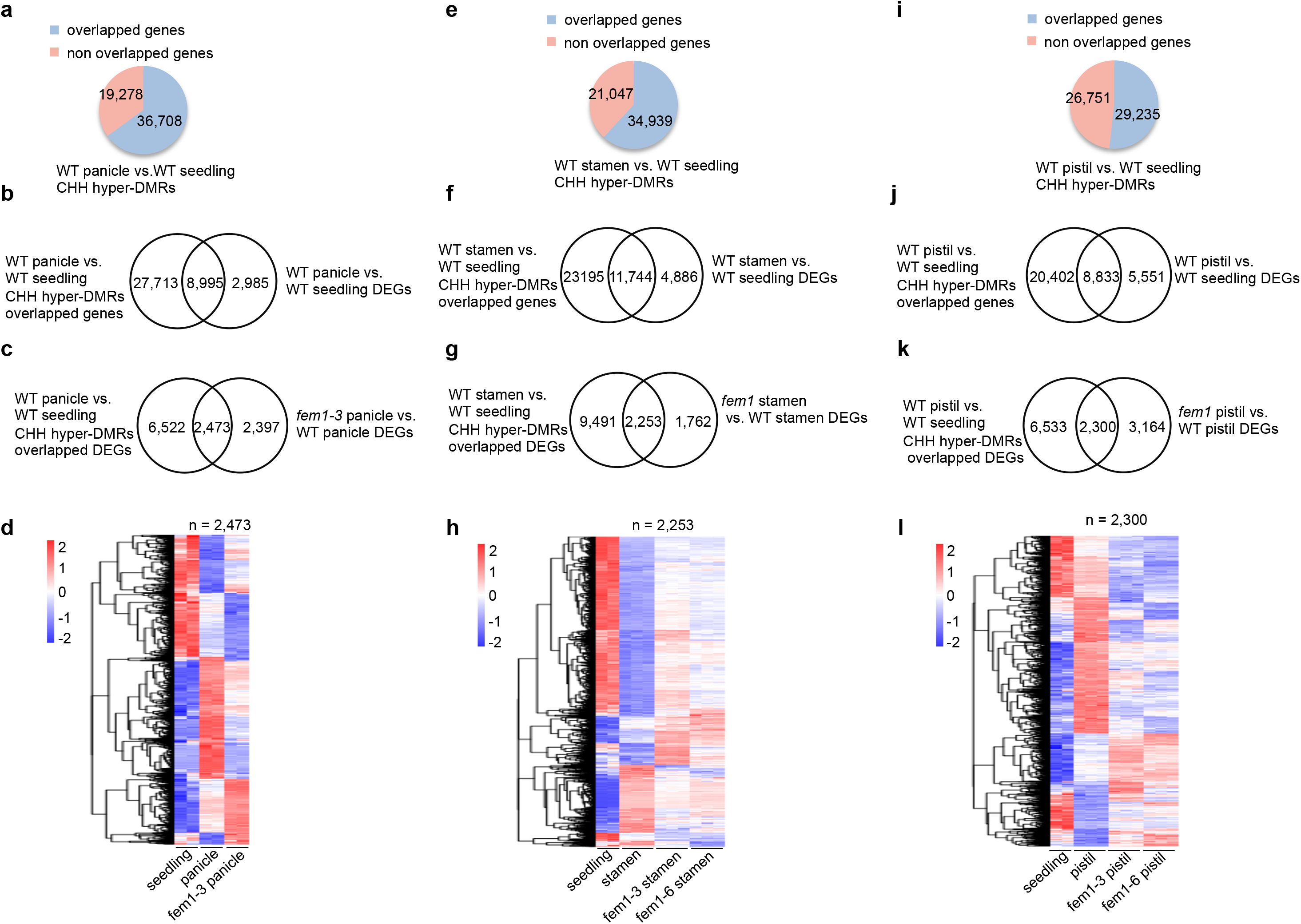
Increase of DNA methylation regulates gene expression. (A) Number of overlapping genes with CHH hyper-DMRs of panicles > seedlings. (B) Venn diagram showing the overlap between genes adjacent to CHH hyper-DMRs of panicles > seedlings and DEGs between panicles and seedlings. (C) Venn diagram showing the overlap between DEGs (WT panicles vs. WT seedlings) overlapped with hyper-DMRs of panicles > seedlings and DEGs (*fem1-3* panicles vs. WT panicles). (D) Heatmap showing the expression levels of 2,473 genes in WT seedlings, WT panicles, and *fem1-3* panicles. (E) Number of overlapping genes with CHH hyper-DMRs of stamens > seedlings. (F) Venn diagram showing the overlap between genes adjacent to CHH hyper-DMRs of stamens > seedlings and DEGs (WT stamens and WT seedlings). (G) Venn diagram showing the overlap between DEGs (WT stamens vs.WT seedlings) overlapped with hyper-DMRs of stamens > seedlings and DEGs (*fem1-3* stamens vs. WT stamens). (H) Heatmap showing the expression levels of 2,253 genes in WT seedlings, WT stamens, and *fem1* stamens. (I) Number of overlapping genes with CHH hyper-DMRs of pistils > seedlings. (J) Venn diagram showing the overlap between genes adjacent to CHH hyper-DMRs of pistils > seedlings and DEGs (WT pistils and WT seedlings). (K) Venn diagram showing the overlap between DEGs (pistils vs. seedlings) overlapped with hyper-DMRs of pistils > seedlings and DEGs between *fem1-3* and WT in pistils. (L) Heatmap showing the expression levels of 2,253 genes in WT seedlings, WT pistils, and *fem1* pistils.

As was the case for panicles and stamens, CHH levels on genes and TEs in WT pistils were higher than the levels on corresponding genes and TEs in WT seedlings (Figures 4D and 4E), and the increase in methylation occurred on short TEs (Supplemental Figure 5H). We identified a total of 70,773 CHH hyper-DMRs of WT pistils vs. WT seedlings (Supplemental Data Set 4) with a total length of 12.2 Mb (Supplemental Figures 5I and 5J), and 59.6% of them were located on TEs (Supplemental Figure 5K). When *fem1-3* pistils were compared with WT pistils, 156,302 CHH hypo-DMRs were identified (Supplemental Figure 5L; Supplemental Data Set 4); these CHH hypo-DMRs covered 12.4% of the rice genome (46.3 Mb, Supplemental Figure 5M), and 58.6% of them were located on TEs (Supplemental Figure 5N). Among the pistil hyper-DMRs, 98.0% overlapped with hypo-DMRs in *fem1-3* pistils (Figure 4F), suggesting that the increase in CHH methylation was largely *FEM1*-dependent. On the specific *FEM1*-dependent DMRs, the CHH methylation level was much higher in WT pistils than in WT seedlings (Figure 4F), suggesting that the increase in methylation might occur on all *FEM1* targets in pistils. The 24-nt siRNA levels, however, were not increased on all *FEM1*-dependent DMRs (Figure 4F).

The CHH hyper-DMRs were longest in panicles (Supplemental Figure 4D), shortest in pistils (Supplemental Figure 5J), and of intermediate length in stamens (Supplemental Figure 5C). However, the three types of DMRs largely overlapped with each other (Supplemental Figure 5O). On seven groups of DMRs, the CHH methylation levels were higher in panicles, stamens, and pistils than in seedlings, and the increase in CHH methylation was reversed in *fem1* mutants of various organs (Supplemental Figure 5P), suggesting that methylation reinforcement largely occurred on common regions in panicles, stamens, and pistils, and that this reinforcement was *FEM1*-dependent.

### Global increase of DNA methylation regulates gene expression in sexual organs

To determine whether the global increase of DNA methylation regulates gene expression in WT panicles, we checked genes near CHH hyper-DMRs (DMRs overlapped with genes from −2 kb in the promoter through 500 bp in the terminator). We found that panicle CHH hyper-DMRs overlapped with 65.6% of all rice genes (36,708 genes, Figure 5A). To investigate the effects of CHH methylation on gene expression, we performed mRNA-seq for WT and *fem1-6* seedlings and for WT, *fem1-6*, and *fem1-3* panicles (Supplemental Data Set 5). Among the 11,980 differentially expressed genes (DEGs, fold-change > 1.2, FDR < 0.05) between panicles and seedlings (Supplemental Data Set 5), 8,995 genes overlapped with panicle hyper-DMRs (Figure 5B), indicating that those gene might be regulated by DNA methylation. In addition, among the 8,995 genes, 2,473 exhibited misregulated expression in *fem1* panicles (Figures 5C and 5D). Among the 49 panicle developmental genes (Supplemental Data Set 6), 35 overlapped with panicle hyper-DMRs (Supplemental Figure 6A). For example, two genes that regulate panicle development, *OsCEP6.1*, which encodes a peptide hormone (Sui et al., 2016), and *OsIAGLU*, which encodes an IAA-glucose synthase (Choi et al., 2012), showed hypermethylation on their promoters in panicles relative to the methylation levels on their promoters in seedlings (Supplemental Figure 6B). The genes were expressed at differential levels in panicles vs. seedlings, and this difference in expression was changed in *fem1* panicles (Supplemental Figure 6B). The misregulation of genes controlling panicle development probably explains the small panicles in the *fem1* mutants (Supplemental Figures 2E and 2F).

**Figure 6.**
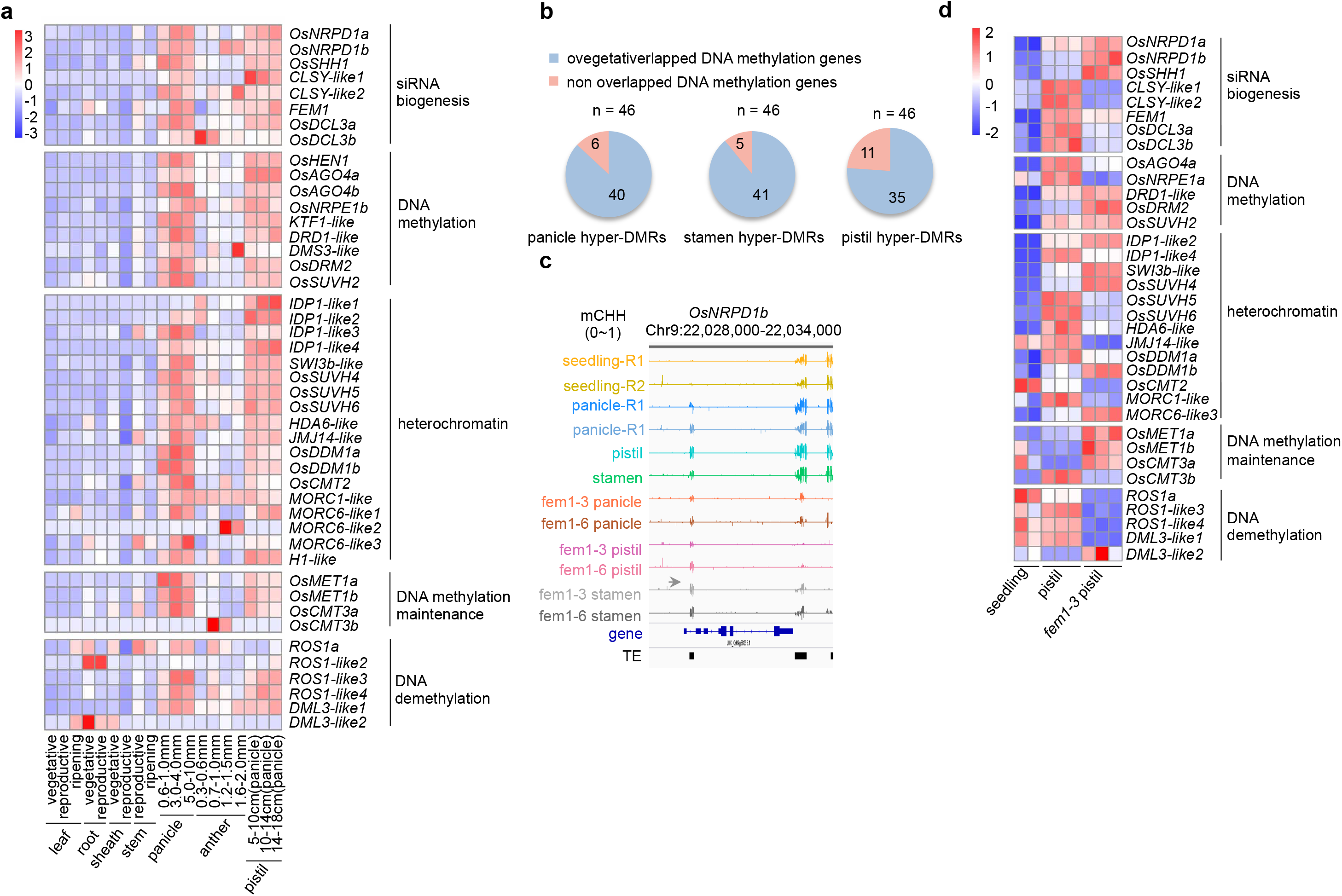
Expression levels of RdDM genes are upregulated in reproductive tissues. (A) Heatmap showing the relative expression levels of DNA methylation-related genes from the Rice XPro database (https://ricexpro.dna.affrc.go.jp/GGEP/index.php). (B) Number of DNA methylation-related genes that overlapped with CHH hyper-DMRs in panicles, stamens, or pistils. (C) Integrated Genome Browser view of CHH methylation levels on and near *OsNRPD1b* in various organs of the indicated genotypes. (D) Heatmap showing the expression levels of DNA methylation-related genes in WT seedlings, WT pistils, and *fem1-3* pistils.

We also performed mRNA-seq for stamens (stage 12) of the WT, *fem1-3*, and *fem1-6* (Supplemental Data Set 1, and Supplemental Data Set 5). Stamen CHH hyper-DMRs overlapped with 62.4% of all rice genes (34,939 genes, Figure 5E). Among the 34,939 genes near DMRs, 11,744 exhibited differential expression between stamens and seedlings (Figure 5F). Of the 11,744 genes, the expression of 2,253 was misregulated in *fem1* stamens (Figures 5G and 5H), suggesting that gene expression in stamens is substantially regulated by hyper-DMRs. Among the 92 identified stamen developmental genes (Supplemental Data Set 6), 66 overlapped with stamen hyper-DMRs (Supplemental Figure 6C). On the border of *Defective Tapetum Cell Death 1* (*DTC1*), which is important for tapetum development (Yi et al., 2016), and on *OsGAMyb*, which encodes a GA-inducible transcription factor (Aya et al., 2009), DNA methylation levels were higher in WT stamens than in WT seedlings, and the methylation level was reduced in *fem1* stamens (Supplemental Figure 6D). Consistent with the latter findings, the transcriptional levels of *DTC1* and *OsGAMyb* were greater in WT stamens than in WT seedlings, and the differences in their transcriptional levels were even greater between *fem1* stamens and WT seedlings (Supplemental Figure 6D), indicating that their proper expression depends on their hypermethylation.

Stage FG8 pistils of the WT, *fem1-3*, and *fem1-6* were also subjected to mRNA-seq analysis (Supplemental Data Set 1; Supplemental Data Set 5). In the WT, pistil CHH hyper-DMRs overlapped with 52.2% of all rice genes (29,235 genes, Figure 5J); of the 29,235 genes, 8,833 showed differential expression between pistils and seedlings (Figure 5J), indicating that their expression could be regulated by hyper-DMRs. Among the 8,833 genes, 2,300 exhibited differential expression in *fem1* pistils compared to the WT pistils (Figures 5K and 5L), suggesting that DNA methylation probably regulates expression of those genes. Among the 20 identified pistil developmental genes (Supplemental Data Set 6), 18 overlapped with pistil hyper-DMRs (Supplemental Figure 6E). For *OsPID*, which regulates the development of stigma (He et al., 2019; Xu et al., 2019), the increase of methylation on its promoter positively regulated its expression in pistils, and that positive regulation was absent in *fem1-3* pistils, indicating that it depended on *FEM1* (Supplemental Figure 6F). The higher expression of *OsOSD1* in WT pistils than in WT seedlings was also correlated with DNA hypermethylation in WT pistils (Supplemental Figure 6F). The mutation in *FEM1* eliminated the DNA methylation and disrupted its expression pattern; downregulation of *OsOSD1* might be responsible for the abnormal formation of female gametes in the *fem1* mutant (Mieulet et al., 2016; Wang et al., 2019).

### Upregulated expression of RdDM machinery genes is responsible for increased CHH DNA methylation

To investigate the mechanism underlying the global increase of DNA methylation in sexual organs of rice, we examined the expression levels of genes related to DNA methylation (Supplemental Table 1) in seedlings, panicles, pistils, and stamens of the WT. The genes necessary for 24-nt siRNA biosynthesis, including *FEM1*, *OsNRPD1*, *OsSHH1*, *CLSYs*, and *OsDCL3*, were upregulated in panicles relative to seedlings, and their transcript level was even greater in pistils (Supplemental Figure 7A). Among those genes, however, only *CLSY-like 1* expression was increased in stamens (Supplemental Figure 7A). Except the genes related to siRNA biosynthesis mentioned above, many of the RdDM pathway genes involved in the methylation of DNA methylation and heterochromatin formation, including *OsNRPE1b*, *OsAGO4*, *OsDMS3*, *OsDRM2*, and *MORC6-like 1*, were upregulated in panicles and pistils (Supplemental Figure 7A). To confirm that most of RdDM genes were upregulated in the sexual organs of rice, we searched the transcriptomes previously reported for rice seedlings, leaves, shoots, panicles, stamens, and pistils (http://rice.plantbiology.msu.edu/expression.shtml). The search indicated that the expression levels of most RdDM genes were higher in panicles and pistils than in leaves, seedlings, or shoots (Supplemental Figure 7B). Moreover, the expression levels of those genes were much higher in young panicles than in mature panicles (Supplemental Figure 7B) but did not differ in young vs. mature pistils (Figure 6A), suggesting that the RdDM machinery was strengthened early in the development of panicles, but was only maintained in pistils during later developmental stages. Another independent search also demonstrated that most RdDM genes in rice are expressed at higher levels in young panicles and pistils than in leaves, roots, sheaths, or stems (Figure 6A, https://ricexpro.dna.affrc.go.jp/GGEP/index.php). Overall, these results indicate that the genes encoding RdDM components are upregulated in panicles and pistils, which reinforces the global DNA methylation in sexual organs.

To determine the molecular mechanism underlying the upregulation of RdDM genes in sexual organs of rice, we examined the DNA methylation levels on 46 DNA methylation-related genes in different organs (Supplemental Table 1), and found that the majority of those genes overlapped with three types of hyper-DMRs (40 genes with panicle hyper-DMRs, 41 genes with stamen hyper-DMRs, and 35 genes with pistil hyper-DMRs, Figure 6B). For example, DNA methylation levels on *FEM1*, *OsDRM2*, and *OsNRPD1b* were greater in panicles, stamens, and pistils than in seedlings (Supplemental Figure 7C; Figure 6C). The high methylation levels on those regions were reduced in panicles, stamen, and pistils of *fem1* mutants (Supplemental Figure 7C; Figure 6C), suggesting that the increase in the methylation of those genes was *FEM1*-dependent. The misregulation of RdDM genes in *fem1* pistils and *fem1* panicles indicated a positive feedback between a global increase in DNA methylation and the up-regulation of the RdDM machinery in the sexual organs of rice (Supplemental Figure 7D; Figure 6D).

## DISCUSSION

DNA methylation is an important epigenetic mark that controls the transcription of genes and TEs, and is essential for genome stability. In mammals including humans, cytosine DNA methylation mainly occurs in the CG context, and reprogramming of DNA methylation is important for gamete and embryo development. Mutation in genes controlling DNA methylation and DNA demethylation in mammals often results in death (Smith and Meissner, 2013; Greenberg and Bourc His, 2019). In Arabidopsis, a dicotyledonous plant with a small genome, most DNA-methylation mutants can reproduce normally, and the defects in *de novo* methylation via the RdDM pathway do not result in any obvious developmental abnormalities (Matzke and Mosher, 2014). These findings with Arabidopsis suggest that *de novo* DNA methylation may not be crucial for plant development. The results presented here, however, indicate that *de novo* DNA methylation is crucial for the reproductive development of rice plants.

In the current investigation of rice, a monocotyledonous plant that diverged 150 millions years ago and is more representative of higher plants than Arabidopsis is in terms of genome size and TE content, we found that mutation in *FEM1* greatly reduced *de novo* DNA methylation and caused total sterility because of developmental defects in both male and female reproductive organs. We demonstrated that the upregulation of RdDM activity including FEM1 activity is responsible for the global increases in DNA methylation in sexual organs of rice. These results therefore indicate that *de novo* DNA methylation through enhancement of RdDM is essential for reproductive development of rice plants.

In Arabidopsis, hypomethylation in the vegetative cells releases the expression of many TEs in the pollen grains (Slotkin et al., 2009). The high methylation level in stamens might make active or inactive DNA demethylation possible in the vegetative cell of Arabidopsis pollen grains. The hypomethylation in TEs in the Arabidopsis vegetative cell might cause the production of easiRNAs (epigenetic activated siRNAs), which were transferred to the sperm cell to repress TE expression (Slotkin et al., 2009; Calarco et al., 2012; Kawashima and Berger, 2014; Kim et al., 2019). The paternal easiRNAs of Arabidopsis also function in post-fertilization genome stability and seed viability (Borges et al., 2018; Martinez et al., 2018). The hypomethylation in a locus-specific manner through DNA demethylation in central cells is inherited in the endosperm and results in gene imprinting in rice and Arabidopsis (Park et al., 2016). For both species, the difference in DNA methylation in different cells of the gametophyte is essential for genome stability and seed viability. Based on the current results, we speculate that the increase in DNA methylation mediated by RdDM in reproductive organs is essential for establishing a specific methylome in various cells of the rice gametophyte. The role of RdDM during meiosis and mitosis in gamete development of rice warrants additional research.

With the global increase in methylation in panicles, stamens, and pistils of rice plants, genes encoding RdDM components were upregulated in panicles and pistils but not in stamens. The reason for this inconsistency between DNA methylation and expression of RdDM genes in stamens is unclear. It is possible that, at an early stage of stamen development, genes in the RdDM pathway are upregulated and that their protein level but not their rates of transcription are maintained at high levels, which would ensure high CHH methylation in stamens at later stages of development.

Although *FEM1* and *OsNRPD1* were upregulated in the sexual organs of rice, the abundance of 24-nt siRNAs was not increased on those developmental hyper-DMRs. The lack of correlation between 24-nt siRNA abundance and DNA methylation levels indicates that 24-nt siRNAs are not sufficient or perhaps even necessary to catalyze DNA methylation in the RdDM pathway. This possibility mirrored our previous finding that following mutations of all four Dicers, substantial DNA methylation remained on RdDM loci in Arabidopsis (Yang et al., 2016). The function of siRNAs in the RdDM machinery requires additional study in rice.

## MATERIALS AND METHODS

### Plant materials

*OsGA2ox1*-overexpression (GAE), *OsGA2ox1*-silencing (GAS), and *fem1* mutants (*fem1-1* and *fem1-2*) are in the background of Taipei 309 (TP309) (*Oryza sativa*, Geng/japonica). The *fem1* mutants (*fem1-3*, *fem1-4*, *fem1-5*, and *fem1-6*) were created using CRISPR/Cas9 technology with the pCBSG03 vector (Zhang et al., 2014) in the background of Nipponbare (*Oryza sativa*, Geng/japonica), which was the WT in this study.

Two kinds of panicles (panicle-R1 and panicle-*fem1-3*; R1 indicates the WT-Replicate 1) that had grown in a field in Hainan were harvested at anther developmental stage 12 in April of 2016. Two other kinds of panicles (panicle-R2 and panicle-*fem1-6*; R2 indicates the WT-Replicate 2) that had grown in a field in Nanjing were harvested at the same developmental stage in September of 2018. Mature stamens and pistils of the WT and mutants (*fem1-3* and *fem1-6*) were harvested in Nanjing in August of 2019. The DNA and RNA isolated from panicles were used for high throughput-sequencing. Eighteen-day-old seedlings without roots (seedling-R1, seedling-R2, and *fem1-6-*seedling; R indicates WT-Replicate) grown in Kimura B nutrient solution in a greenhouse (16 h/8 h light/dark with a 28℃/25℃ cycle) were collected, and their DNA and RNA were isolated for high-throughput sequencing.

### Genetic screen of *five elements mountain* mutants

GAS seeds (about 50,000) were soaked in water for 24 h at room temperature and then transferred into 1% EMS (ethyl-methanesulfonate) for another 24 h. The EMS-treated seeds were planted in an isolated paddy field. The seeds were separately harvested from each individual plant (more than 23,000 M1 plants). The M2 plants from the same M1 line were planted together. The M2 plants that produced dwarf seedlings were selected and planted (about 200 M1 lines). When the dwarf plants were 3 months old, their *OsGA2ox1* transcript levels and those of GAS plants were assessed. Those dwarf rice plants with *OsGA2ox1* transcript levels that were at least 20 times greater than those of GAS were named *fem*. The name was inspired by the journey to the west, in which Buddha created Five Elements Mountain (‘FEM’) to imprison a monkey king who disturbed the hierarchy in heaven.

### Map-based cloning of *FEM1*

To clone *FEM1*, heterozygous *fem1* plants were crossed with TN1 plants (cv. Taichung Native 1), a Xian/indica variety. Seeds of individual F1 plants were separately collected and planted. The dwarf seedlings from the F2 population were selected and served as mapping samples. Using 228 SSR and STS molecular markers, we conducted bulked segregant analysis (BSA) of mutant and WT pools (mutant and WT pools consisted of 30 dwarf plants or normal plants, respectively). R1 and R7 were first identified as linked markers in BSA. Fine mapping was then conducted using 140 dwarf plants to narrow the location of *fem1-1* between R4 and R5. Finally, a large mapping population consisting of 890 samples was used to determine the region that contained *fem1-1* to 103 kb. A similar procedure was used to map *fem1-2* (Supplemental Figure 1I). The oligos of molecular markers used for fine mapping are listed in Supplemental Table 2.

### Vector construction and plant transformation

To generate the *OsGA2ox1* overexpression construct, the 8,515-bp fragment containing the *OsGA2ox1* coding region and its promoter and terminator was isolated from BAC OsJNBa0017J22, which was digested by restriction enzyme *Xba*I (NEB, R0145S). With the T4 DNA ligase (TAKARA, 2011A), the product was ligated to the binary vector pCAMBIA1301-35SN, which was digested by *Xba*I (NEB, R0145S). The construct with the desired insertion was isolated by colony-PCR (Supplemental Table 2). Agrobacterium EHA105 with the sequenced vector was used for rice transformation.

For the complementary construct, an 8,339-bp DNA fragment composed of the coding region, the 2,098-bp promoter, and the 1,030-bp terminator of *FEM1* was amplified using high fidelity polymerase (Vazyme, P505-d1) and the oligos listed in Supplemental Table 2. The PCR product was cloned into the binary vector pCAMBIA3301, which was digested by *Eco*RI and *Hind*III (NEB, R3101S and R3104S) using the ClonExpress MultiS One Step Cloning Kit (Vazyme, C113-02) according to the manufacturer’s instructions. Agrobacterium EHA105 with the sequenced vector was used for rice transformation.

Two sgRNAs targeting *FEM1* were designed on http://skl.scau.edu.cn/, and the oligos were synthesized in Genescript (Supplemental Table 2). After gradient annealing, the two oligos were cloned into pCBSG03, which was digested by *Bsa*I (NEB, R0535S) as previously reported (Zhang et al., 2014). Agrobacterium EHA105 with the sequenced vector was used for rice transformation.

### Pollen viability assay

Mature anthers of various genotypes were fixed in 50% FAA (5% formaldehyde, 5% glacial acetic acid, and 63% ethanol) for 24 h. The pollen was then released in a drop of 1% I_2_-KI on a glass slide. After they were stained for 3 min at room temperature, the pollen grains were examined with a light microscope (Olympus, BX53).

### Semi-thin sections

Anthers of *fem1* and the WT at various stages (Zhang et al., 2011) were fixed in 50% FAA, and samples were successively dehydrated in 50% ethanol for 30 min, 70% ethanol for 30 min, 70% acetone for 20 min, 80% acetone for 20 min, 90% acetone for 20 min, 95% acetone for 30 min, and finally three times in 100% acetone for 40 min each time. The anthers were then placed in 3:1 acetone: embedding medium (Zhongjingkeyi Technology Co., Ltd. SPI-Pon 812 kit, GP18010) for 2 h, 1:1 acetone: embedding medium for 4 h, and 1:3 acetone: embedding medium for 12 h. The anthers were finally kept at 65℃ for 48 h in embedding medium. Microtome sections (2 μm thick) were stained with 0.05% toluidine blue and observed with a light microscope (Olympus, BX53).

### Scanning electron microscopy (SEM) of pollen

For SEM of pollen, mature flowers (anthers at stage 12) were fixed in 2.5% glutaraldehyde overnight at 4℃ and then dehydrated in an ethanol series of 30, 50, 70, 80, 90, and 100% (5 min for each step). The samples were then critical point dried and coated with gold powder. Images were obtained with a HITACHI SU80I0 scanning electron microscope.

### Observation of embryo sacs

Embryo sacs were stained and observed as previously described with slight modification (Zhao et al., 2013). In brief, mature florets of the WT and *fem1* mutants were collected and immediately fixed in 50% FAA for 24 h at room temperature. The samples were then stored in 70% ethanol at 4℃. Before staining, the ovaries were dissected with the aid of a stereomicroscope. The tissue was processed through an ethanol series (50, 30, and 15%, 30 min for each concentration) and finally transferred into distilled water for 30 min. The ovaries were successively stained in 2% aluminium potassium sulfate for 20 min, 4% eosin for 12 h, and 2% aluminium potassium sulfate for 20 min; they were then washed with distilled water for 24 h, followed by three washes with distilled water. The tissue was dehydrated in an ethanol series (30, 50, 70, 80, 90, and 100%, 30 min for each step, but three times in 100% ethanol). The ovaries were then placed in 1:1 ethanol:methylsalicylate for more than 1 h, and cleared in methylsalicylate three times for 1 h each time. The embryo sacs were examined with a confocal laser scanning microscope (Zeiss LSM780) at 514 nm wavelength.

### RT-PCR

Total RNA was extracted using TRIzol reagent (Invitrogen, 15596018). About 1 μg of total RNA was used to synthesize cDNA with the HiScript® II Q RT SuperMix kit (Vazyme Biotech Co., Ltd. R223-01) according to manufacturer’s protocol. The *OsActin1* gene (LOC_Os03g50885) was used as the internal control. The gene-specific oligos used for RT-PCR are listed in Supplemental Table 2.

### Chop-PCR assay

A 200-ng quantity of genomic DNA was digested by the methylation-sensitive enzyme *Hae*III (NEB, R0108S) in a 20-μl reaction mixture at 37℃ for 20 h. PCR was performed using 2 μl of the digested DNA as template in a 20-μl reaction mixture with the locus-specific oligos (Supplemental Table 2).

### Whole-genome bisulfite sequencing

Genomic DNA was treated with the EZ DNA Methylation-Gold kit (ZYMO, D5005). The libraries were prepared according to NGS Fast DNA Library Prep Set for Illumina protocols, and were sequenced on the Hiseq-Xten platform. The reads were aligned to the converted rice reference genome of release 7 (Kawahara et al., 2013) using bismark (Krueger and Andrews, 2011) (version 0.18.1) with default parameters. The reads uniquely mapped to the chloroplast genome were used to calculate the conversion rate. The methylation level of each cytosine site or region was calculated as the total number of methylated cytosines divided by the total number of sequenced cytosines of the site or region.

The average methylation levels of CG, CHG, and CHH on genes and TE regions, and in their 3-kb upstream and downstream 3-kb regions, were analyzed in windows of 100 bp. For the average methylation levels of CG, CHG, and CHH on TEs of different length, each TE surrounding region (2-kb regions upstream and downstream) was divided into 20 windows (100 bp in length), and 20 windows of the TE region were equally divided according to TE length.

### DMR identification and analysis

For identification of DMRs, only cytosine covered by at least four reads was considered an effective site. DMRs were searched using a 100-bp sliding window with a 50-bp step-size. Windows with at least four effective cytosines were retained. DNA methylation levels were compared pairwise with Fisher’s exact test, and the *p*-values were adjusted for multiple comparisons using the Benjamini-Hochberg method. Windows with FDR < 0.01 were selected. Finally, regions with an absolute methylation level difference of 0.4, 0.2, and 0.1 for CG, CHG, and CHH, respectively, were retained for subsequent analysis (Stroud et al., 2013). Neighboring DMRs were merged if the gap was < 100 bp. All of the analyses were conducted with R (version 3.1.0). The overlaps between DMRs were calculated with bedtools (version 2.25.0).

### mRNA sequencing and analysis

RNA-seq libraries were constructed with the NEBNext Ultra RNA Library Prep Kit of Illumina (NEB, E7530L), and were then sequenced on the Hiseq-Xten platform. The low-quality reads in the raw data were removed by Trimmomatic (version 0.33), and the clean reads were then aligned to the rice reference genome of release 7 (Kawahara et al., 2013) using TopHat-2.1.1 (Trapnell et al., 2012). The expression level for each gene was calculated using the FRKM (fragments per kilobase of exon per million mapped reads) method. Cuffdiff (version 2.2.1) was used for differential expression analysis (Trapnell et al., 2009; Trapnell et al., 2012). The DEGs between two samples were selected using the following criteria: the fold-change (FPKM values among replicates were averaged) was > 1.2, and the FDR was < 0.05. Heatmaps of gene expression levels were generated by R (version 3.1.0).

### Small RNA sequencing and analysis

After total RNAs were isolated, the 18 to 45 nt small RNAs were cut from the PAGE gel and then ligated to adapters. The adapted small RNAs were reversely transcribed into cDNA. PCR was used to amplify the cDNA fragments. The PCR products between 110-130 bp were selected and then purified with the QIAquick Gel Extraction Kit (QIAGEN, 28704). Finally, the products were sequenced on the BGISEQ-500 platform. The reads with N were removed from the raw data by script. The clean sRNA reads were then aligned to the rice reference genome of release 7 (Kawahara et al., 2013) using BOWTIE software (Langmead, 2010) (version 1.1.2) with 0 mismatches. The reads that mapped to structural RNA were removed, and the remaining and unique mapped 24-nt reads were used for subsequent analysis. The expression levels were normalized using the reads per ten million mapped reads (RPTM).

## Availability of data

Sequencing data have been deposited in the Gene Expression Omnibus (GSE112259, GSE130168, and GSE152155).

## Author contributions

Yang DL conceived of the project and designed the experiments; Wang L conducted most of the experiments and performed the bioinformatics analysis; Zeng L conducted the genetic screens; Zheng K, Zhu T, Yin Y, and Xu D contributed analyzing agents and tools; Zhan H assisted with the examination of anther and pollen; Wu Y assisted with the bioinformatics analysis; and Yang DL and Wang L wrote the paper.

## Acknowledgements

This study was supported by the National Natural Science Foundation of China (31671340), the Natural Science Foundation of Jiangsu Province (SBK20170027), the National Key Transformation Program (2016ZX08001002), and the Jiangsu Collaborative Innovation Center for Modern Crop Production to DL Yang. We thank Bin Han for providing the BAC clone. The high-throughput sequencing data were analyzed on the high-performance computing platform of the Bioinformatics Center, Nanjing Agricultural University.

## Supplemental Figures

**Supplemental Figure 1.**
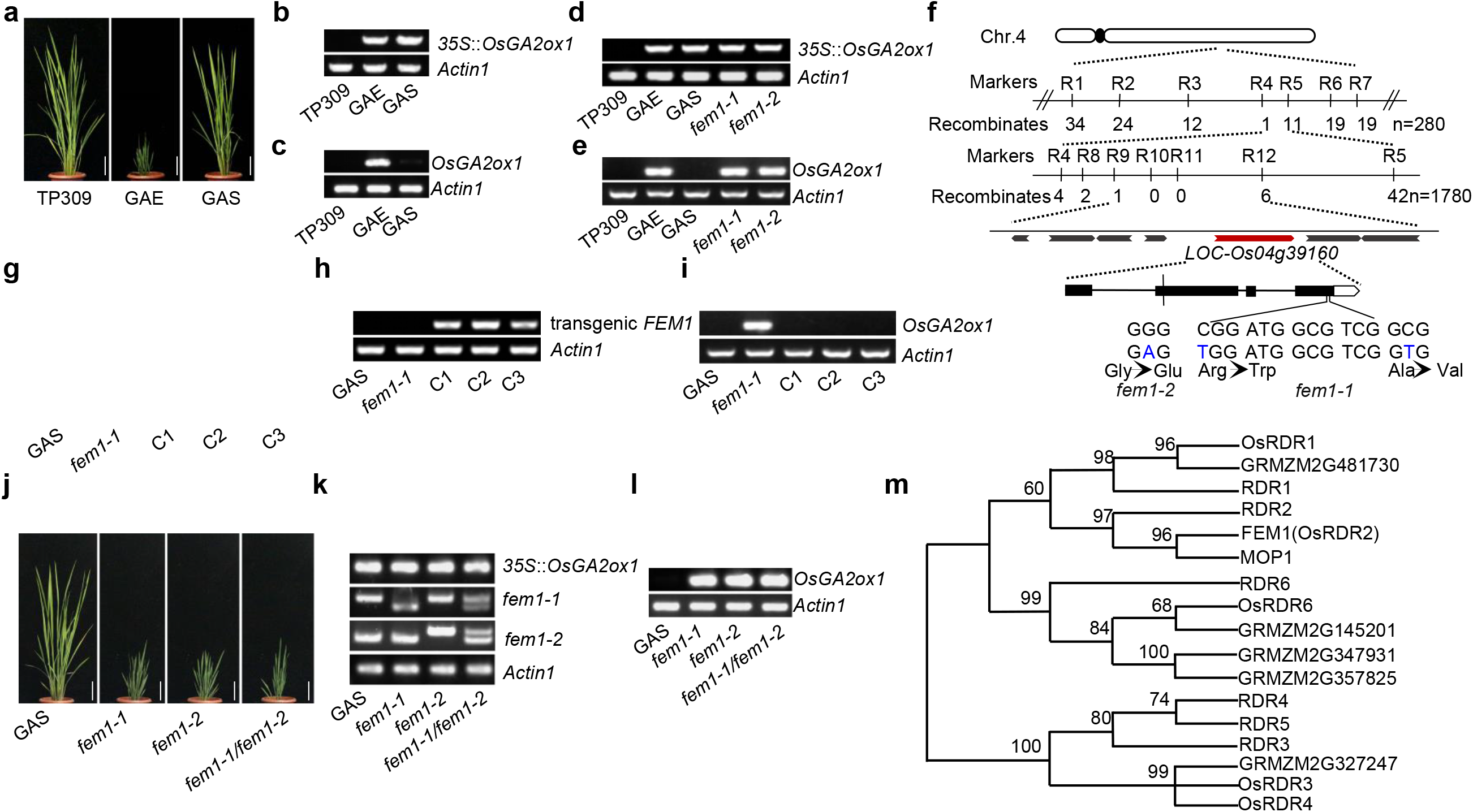
Characterization of the *fem1* mutant and map-based cloning of *FEM1*. (A) Morphology of 80-day-old TP309, GAE, and GAS plants. Scale bar = 10 cm. (B and D) Genotyping transgenic *OsGA2ox1* in TP309, GAE, and GAS. *Actin1* served as the control. (C and E) Transcript level of *OsGA2ox1* as indicated by RT-PCR. *Actin1* served as the control. (F) Diagram showing the map-based cloning of *fem1-1*. The mutations of *fem1-1* and *fem1-2* are indicated. (G) Morphology of 3-month-old plants of GAS, *fem1-1*, and three complementary lines (C1-C3). Scale bar = 10 cm. (H) Genotyping of transgenic *FEM1* in various genotypes. (I) Transcript level of *OsGA2ox1* in the indicated genotypes. (J) The morphology of the indicated genotypes. Scale bar = 10 cm. (K) Genotyping of *fem1-1* and *fem1-2* in the indicated genotypes. (L) Transcript levels of *OsGA2ox1* in the indicated genotypes as determined by RT-PCR. (M) Phylogenetic analysis by MEGA 6.0 of the RDR family from rice, Arabidopsis, and maize.

**Supplemental Figure 2.**
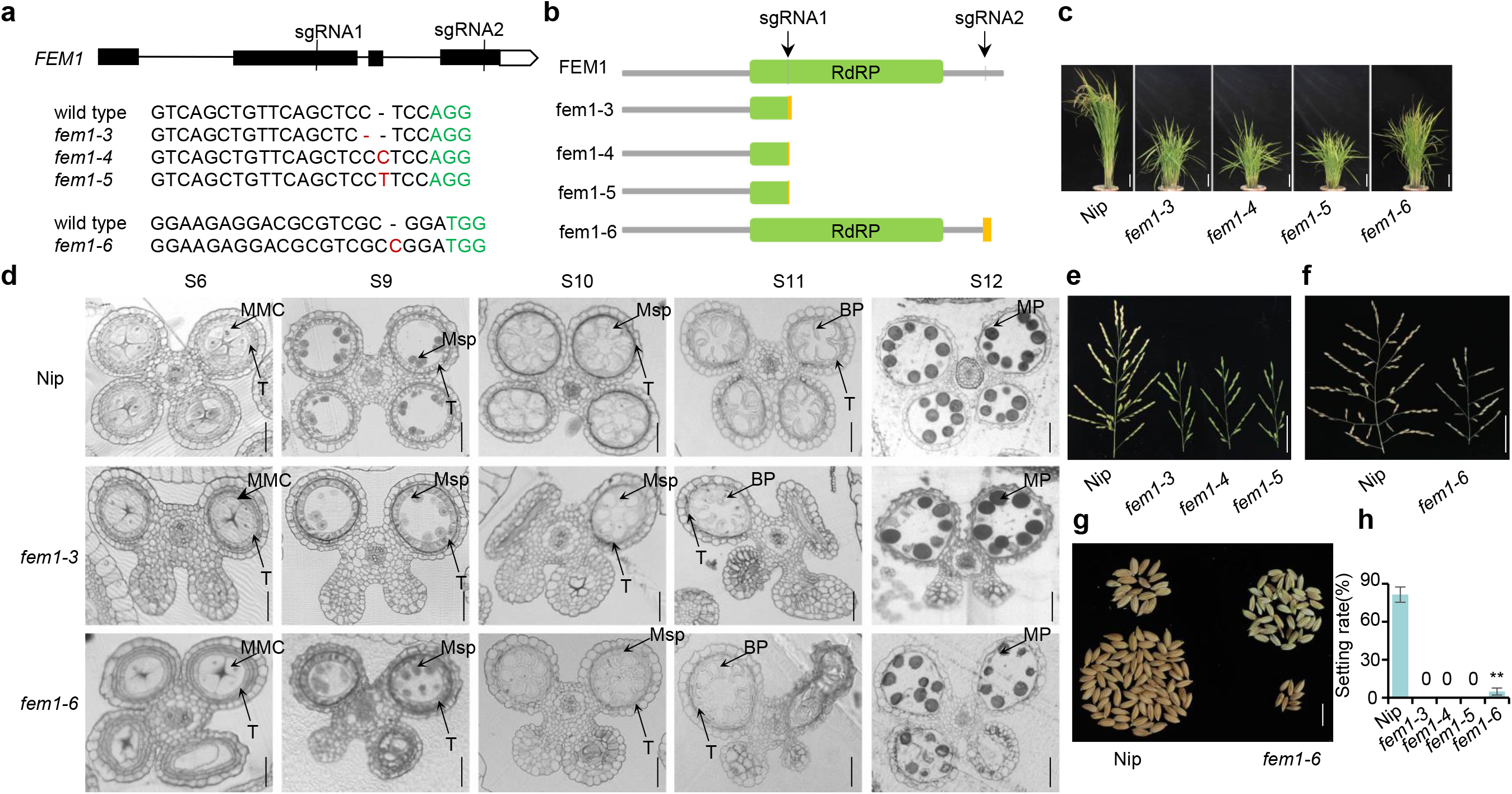
*fem1* exhibits multiple developmental defects. (A) Diagram showing the sites of sgRNA1 and sgRNA2 on *FEM1*. *fem1-3*, *fem1-4*, and *fem1-5* were generated by sgRNA1, and *fem1-6* was generated by sgRNA2. (B) Schematic diagram showing the protein structure of FEM1 and various *fem1* mutants. The green frame indicates the RdRP domain. The yellow box indicates the mutated protein fragment. The arrows indicate the sgRNA position. (C) Morphology of the WT and various *fem1* mutants. Scale bar = 10 cm. (D) Transverse sections of anthers at five stages of development in the indicated genotypes. MMC: microspore mother cell, Msp: microspore, BP: binucleate pollen grain, MP: mature pollen grain, T: tapetum. Scale bar = 10 μm. (E) Panicle morphology of the WT, *fem1-3, fem1-4*, and *fem1-5*. Scale bar = 5 cm. (F) Panicle morphology of the WT and *fem1-6*. Scale bar = 5 cm. (G) Unfilled (top) and filled (bottom) rice grains of the WT and *fem1-6*. Scale bar = 1 cm. (H) Seed setting rate in various genotypes (n = 15).

**Supplemental Figure 3.**
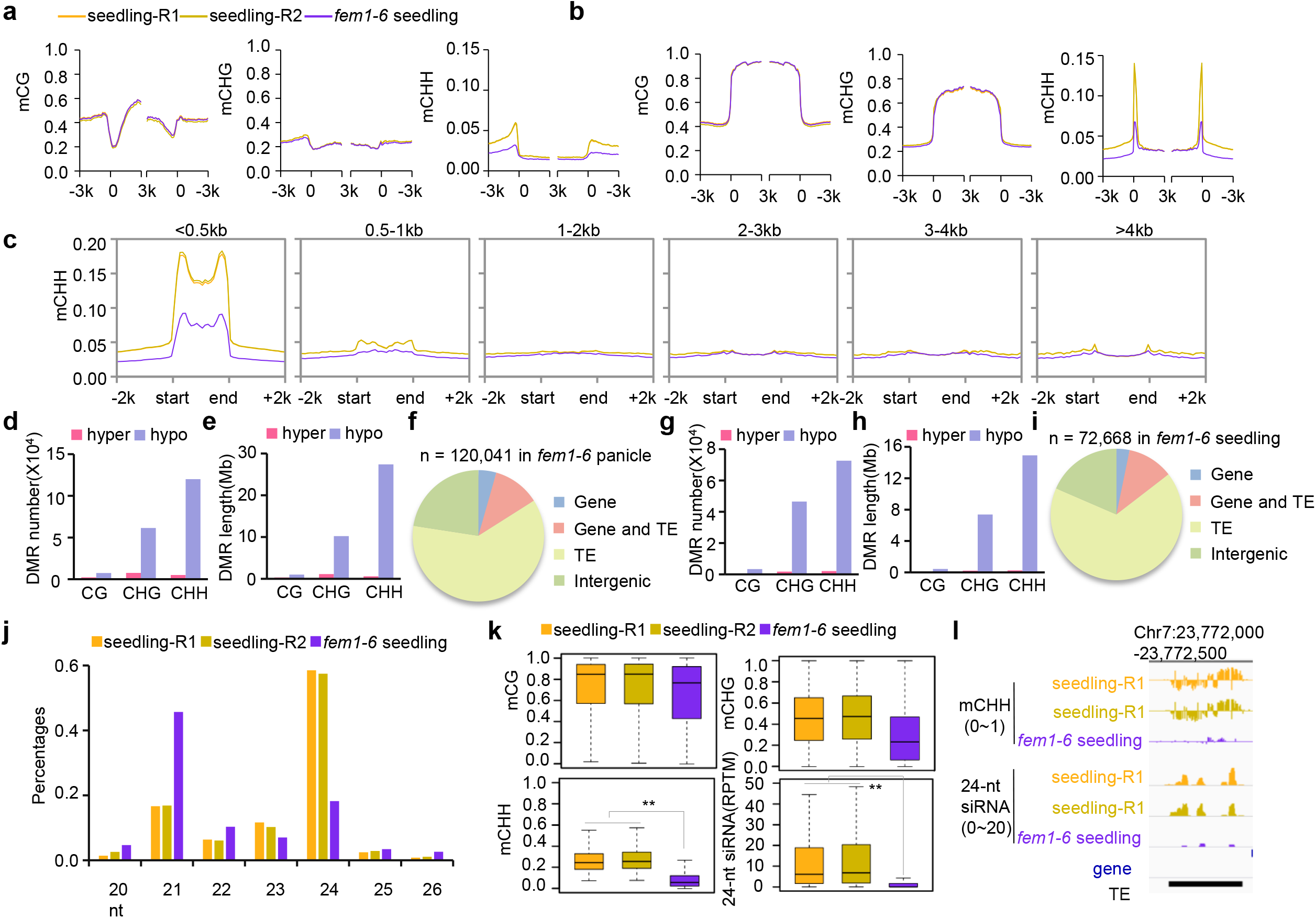
FEM1 is involved in maintaining CHH methylation in panicles and seedlings. Normalized CG, CHG, and CHH methylation levels on genes (A) and TEs (B) in WT and *fem1-6* seedlings. (C) Normalized CHH methylation levels on TEs of different length in WT and *fem1-6* seedlings. Number (D) and length (E) of various DMRs in *fem1-6* panicles compared to WT panicles. (F) Genomic distribution of CHH hypo-DMRs in *fem1-6* panicles. The number (G) and length (H) of various DMRs in *fem1-6* seedlings compared to WT seedlings. (I) Genomic distribution of CHH hypo-DMRs in *fem1-6* seedlings. (J) Abundance of small RNAs with different lengths in WT and *fem1-6* seedlings. (K) Box-plots indicating the methylation level of CG, CHG, and CHH and 24-nt siRNA abundance on CHH hypo-DMRs in *fem1-6* seedlings (**, *p*<0.01 by Wilcoxon sum test). (L) Integrated Genome Browser view of CHH methylation and 24-nt siRNA abundance in one representative CHH hypo-DMR in *fem1-6* seedlings.

**Supplemental Figure 4.**
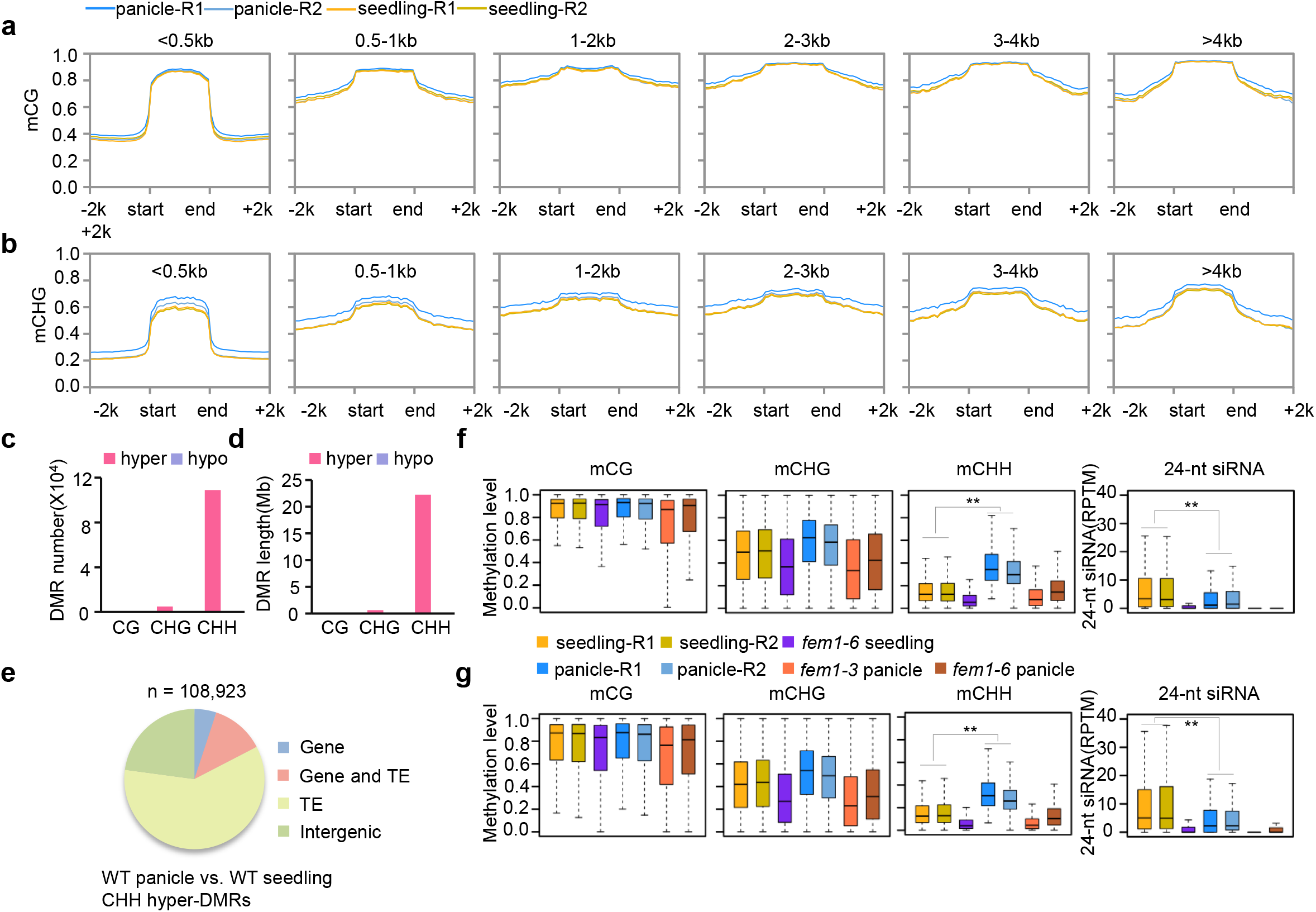
FEM1 is involved in maintaining CHH methylation in panicles. Normalized CG (A) and CHG (B) methylation levels on TEs of different length in panicles and seedlings. Number (C) and length (D) of various DMRs between panicles and seedlings. (E) Genomic distribution of panicle CHH hyper-DMRs. Box-plots showing the methylation level of CG, CHG, and CHH and 24-nt siRNA abundance of the indicated genotype on CHH hyper-DMRs of panicles > seedlings (F), and on CHH-hypo-DMRs of *fem1-3* panicles < WT panicles (G) (**, *p*<0.01 by Wilcoxon sum test).

**Supplemental Figure 5.**
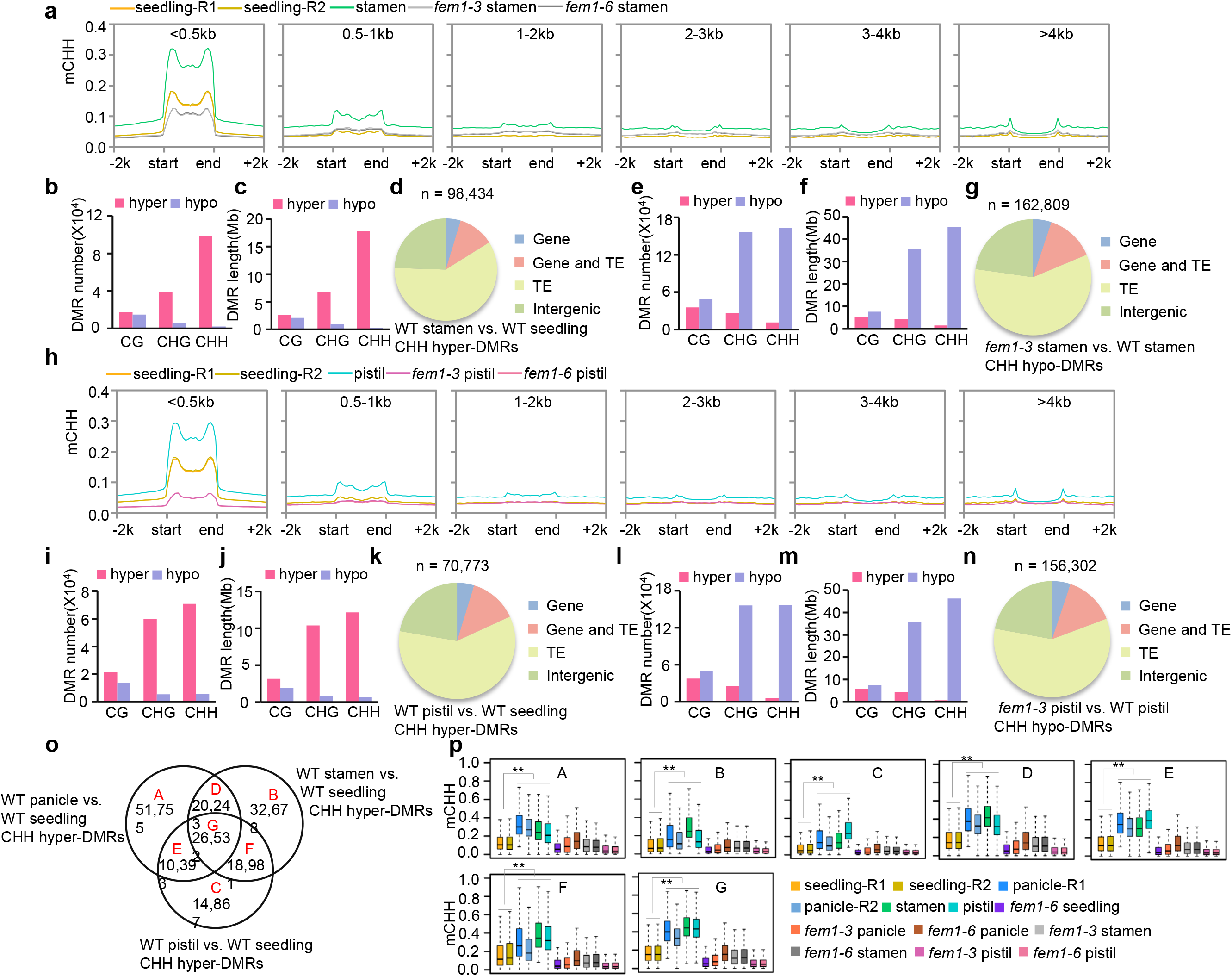
Stamens and pistils are CHH hyper-methylated compared with seedlings. (A) Normalized CHH methylation levels on TEs of different length in seedlings and stamens of different genotypes. The number (B) and length (C) of various DMRs between stamens and seedlings. (D) Genomic distribution of stamen CHH hyper-DMRs. Number (E) and length (F) of various DMRs in *fem1-3* stamens. (G) Genomic distribution of *fem1-3* CHH hypo-DMRs in stamens. (H) Normalized CHH methylation levels on TEs of different length in seedlings and pistils of different genotypes. Number (I) and length (J) of various DMRs between pistils and seedlings. (K) Genomic distribution of pistil CHH hyper-DMRs. Number (L) and length (M) of various DMRs in *fem1-3* pistils. (N) Genomic distribution of CHH hypo-DMRs of *fem1-3* pistils < WT pistils. (O) Venn diagram showing the number of overlapping loci among the CHH hyper-DMRs in panicles, stamens, and pistils. (P) Box-plots showing the CHH methylation level of the various genotypes on different types of DMRs.

**Supplemental Figure 6.**
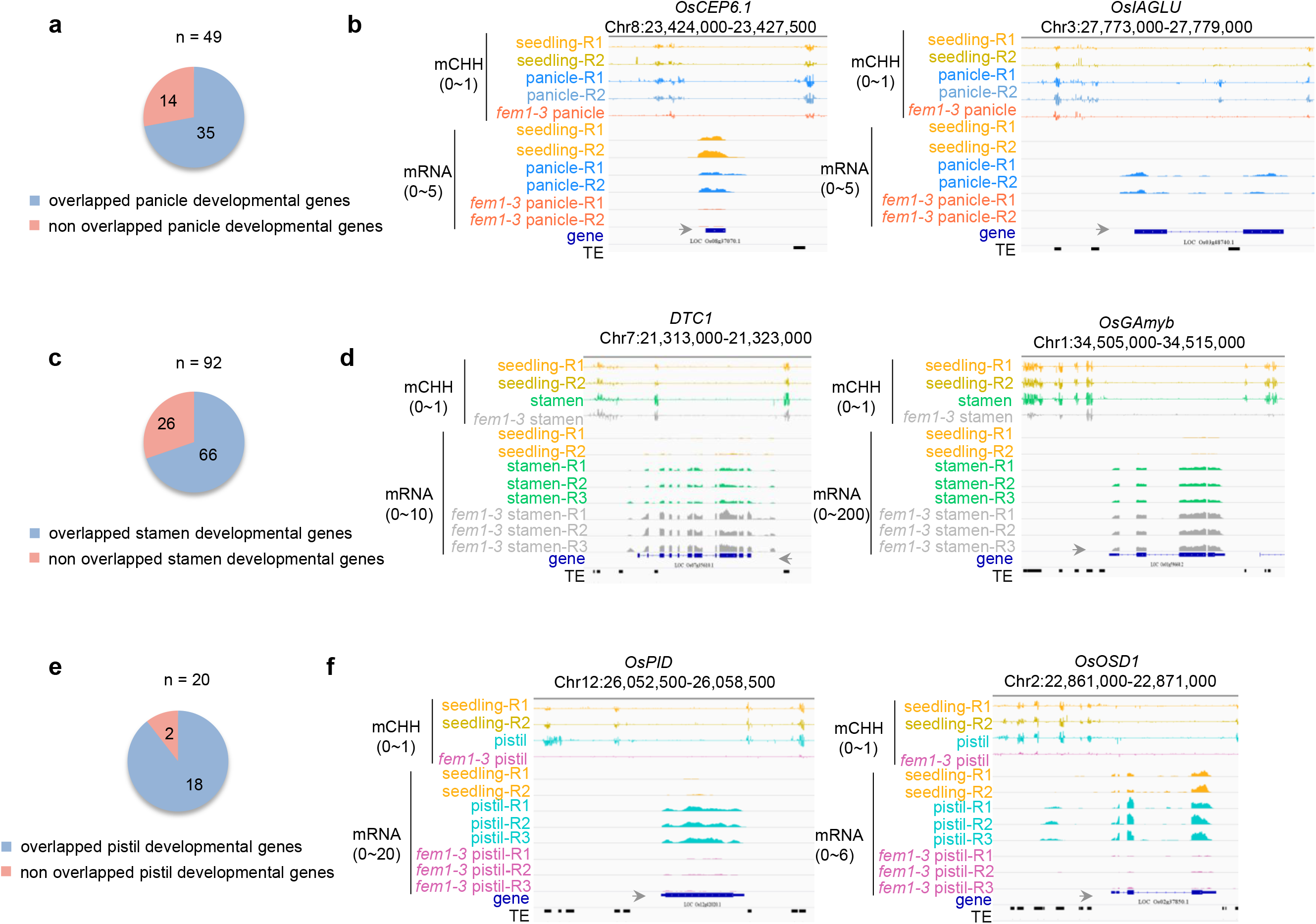
CHH hyper-DMRs regulate reproductive development. (A) Number of genes related to panicle development that overlapped with panicle CHH hyper-DMRs. (B) Integrated Genome Browser view of CHH methylation levels and mRNA levels in WT seedlings, WT panicles and *fem1-3* panicles. (C) Number of genes related to stamen development that overlapped with stamen CHH hyper-DMRs. (D) Integrated Genome Browser view of CHH methylation levels and mRNA levels in WT seedlings, WT stamens and *fem1* stamens. (E) Number of genes related to pistil development that overlapped with pistil CHH hyper-DMRs. (F) Integrated Genome Browser view of CHH methylation levels and mRNA levels in WT seedlings, WT pistils and *fem1* pistils.

**Supplemental Figure 7.**
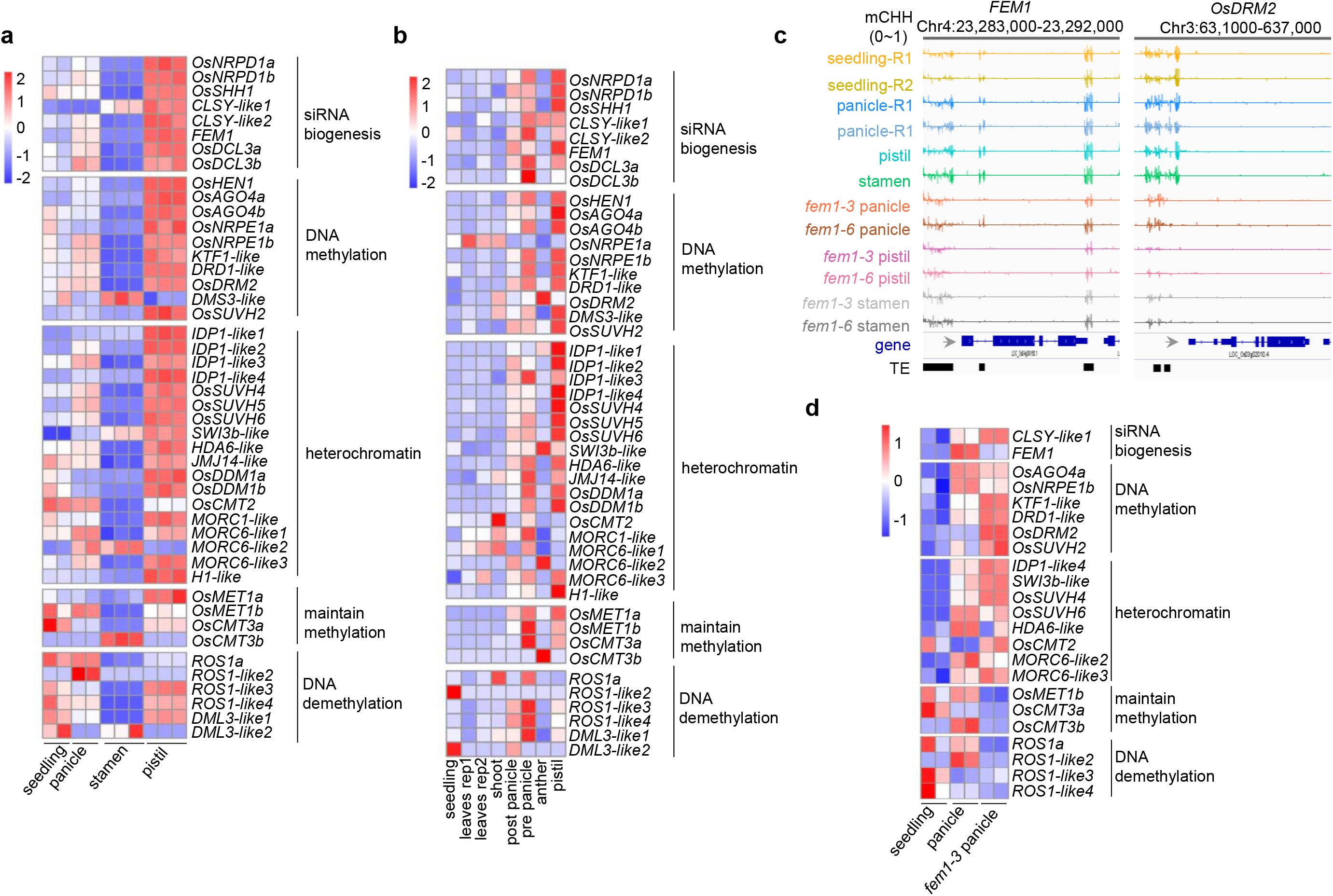
Upregulation of genes related to DNA methylation in reproductive tissues. (A) Expression levels of genes related to DNA methylation in seedlings, panicles, stamens, and pistils as indicated by our mRNA-seq analysis. (B) Expression levels of genes related to DNA methylation in seedlings, leaves, shoots, panicles, anthers, and pistils from the MSU database (http://rice.plantbiology.msu.edu/expression.shtml). (C) Integrated Genome Browser view of CHH methylation levels on *FEM1* and *OsDRM2* in various organs of the indicated genotypes. (D) Heatmap showing the expression level of genes related to DNA methylation in WT seedlings, WT panicles and *fem1-3* panicles.

**Supplemental Data Set 1.** Basic information of high-throughput sequencing.

**Supplemental Data Set 2.** Panicle hyper-DMRs and *fem1* hypo-DMRs in panicles.

**Supplemental Data Set 3.** Stamen hyper-DMRs and *fem1* hypo-DMRs in stamens.

**Supplemental Data Set 4.** Pistil hyper-DMRs and *fem1* hypo-DMRs in pistils.

**Supplemental Data Set 5.** List of DEGs.

**Supplemental Data Set 6.** List of reproductive regulatory genes.

**Supplemental Table 1.** List of DNA methylation-related genes.

**Supplemental Table 2.** Oligo sequences.

